# Airway mucins function as endogenous inhibitors of neutrophil extracellular traps

**DOI:** 10.64898/2026.05.14.719291

**Authors:** Allison Boboltz, Vaidehi Rathi, Gregg A. Duncan

## Abstract

Neutrophils recruited to the airways are important for innate lung defense and can release neutrophil extracellular traps (NETs) to capture and eliminate microbes. While NETs are not abundant in healthy airways, uncontrolled NETosis is a known pathological feature and contributor to both chronic and acute respiratory diseases. Prior studies have shown that mucin glycoproteins secreted in the oral cavity and cervicovaginal tract can modulate NETosis, but it remains unknown whether mucins secreted in the respiratory tract influence NET formation. In these studies, we discovered that human airway mucus strongly inhibits NETosis in primary human neutrophils in a sialic acid dependent manner. In comparison, mucus produced by human airway epithelial cells genetically engineered to lack either MUC5B or MUC5AC secreted airway mucins showed a reduced ability to suppress NETosis. To assess how the lung microenvironment in obstructive lung diseases may influence mucus-dependent NET formation, we engineered a synthetic, mucin-laden hydrogel model with physical properties resembling that of mucus in a healthy lung and a disease-affected lung. When neutrophils were cultured on these gel substrates, we found that increasing gel stiffness led to a significantly greater extent of NETosis. Together these data demonstrate a new functional role of airway mucus in modulating neutrophil homeostasis in the respiratory tract and provide evidence that mucus dysfunction in disease can impair its ability to regulate NETosis.

## Introduction

Gel-forming mucins are continuously secreted in the lung to form a physical barrier to prevent damage to the underlying epithelium. This mucus is cleared from the airways to remove inhaled particles and pathogens via the coordinated beating of ciliated epithelial cells in a process known as mucociliary transport.^1^ One of the primary components of mucus is mucin glycoproteins, with ∼80% of their mass being glycans.^1^ Mucin 5B (MUC5B) and mucin 5AC (MUC5AC) are the two secreted mucin subtypes forming the mucus gel layer in the airways. Resident and infiltrating immune cells present in the lung are also important in eliminating inhaled particles and pathogens to maintain pulmonary health, such as macrophages and neutrophils. In induced sputum samples from healthy individuals, approximately 40-50% of the identified cell population is neutrophils.^2,3^ However, the proportion of neutrophils in the airways often rises dramatically in chronic lung diseases such as asthma, cystic fibrosis (CF), and chronic obstructive pulmonary disease (COPD).^3,4^ Sputum neutrophilia is also observed in acute viral and bacterial respiratory infections.^5,6^ In considering this, the balance of mucociliary and immune-mediated clearance functions are critical to maintaining lung health.

Neutrophils are the most abundant leukocyte in the body and are often considered the first responders of the immune system. Neutrophils mount antimicrobial and pro-inflammatory responses to infection through degranulation, phagocytosis, and release of neutrophil extracellular traps (NETs) generated through a process also known as NETosis.^7^ NETs are web-like complexes consisting of de-condensed chromatin and various granular proteins that can capture and kill pathogens.^8^ However, NETs are considered a “double edged sword” of innate immunity due to their propensity to damage the host tissues through exposure to potent antimicrobial granular proteins and promotion of downstream inflammation.^9^ NETs are released sparingly in healthy airways, as evidenced by the low amounts found in healthy sputum samples.^10–12^ Analysis of sputum samples and mucus plugs from individuals with various chronic lung diseases including asthma, CF and COPD reveals significant upregulation of NETosis.^10–13^ An increase in NETosis is also observed in acute pulmonary infections including COVID-19, influenza, respiratory syncytial virus, and bacterial pneumonia.^6,14–18^ The over-abundance of DNA from NETs, as well as other necrotic cells, in airway secretions is also known to impair mucociliary function which may in effect exacerbate their effects on airway mucus plugging and inflammation.^19–21^ Despite the importance of NETs in lung function and dysfunction, it is unknown if the airway mucus itself contributes to the modulation of NETosis in health, and if alterations to mucus in obstructive lung disease may contribute to the increase in NETs in disease-affected airways.

Prior studies have investigated the potential role of mucins in the oral cavity and female reproductive tract in modulating neutrophil activation and NETosis.^22–24^ Human saliva induces NETosis in human neutrophils via interactions with mucins rich in sialyl-Lewis X glycans.^23^ Conversely, purified mucins from bovine cervical tissues suppressed NETs in bovine neutrophils via the sialic acid glycans.^22^ Though the same mucins are expressed in multiple mucosal tissues, as MUC5B is a prominent salivary mucin and both MUC5B and MUC5AC are found in the cervical and airway mucus, the glycosylation patterns differ.^1,24,25^ Therefore, these studies point to divergent roles of mucins in regulating NETosis to maintain neutrophil homeostasis at different mucosal interfaces.

Motivated by this prior work, we evaluated if secreted airway mucus can directly regulate NETosis using human *in vitro* monoculture and co-culture models. We also evaluated the role of airway-specific mucin subtypes on the modulation of NETosis using human airway epithelial (HAE) cells genetically engineered to achieve knockout of either MUC5B or MUC5AC. In addition, we developed a tunable hydrogel model to investigate how changes to the mucus gel in disease may disrupt the modulation of NETosis. Ultimately, this study uncovers how mucus in the lung functions to maintain neutrophil homeostasis in healthy airways. Given the contribution of NETs to lung pathology, we also provide new evidence which may explain how physical alterations to mucus can lead to enhanced neutrophilic inflammation in respiratory disease-states.

## Results

### Airway mucus suppresses NETosis

Mucus was collected from human airway epithelial (HAE) cell cultures. We used BCi-NS1.1 cells, an immortalized airway epithelial basal cell line originating from a non-smoking healthy donor.^26^ Importantly, BCi-NS1.1 cells can be differentiated into a functional mucociliated airway epithelium when cultured at the air-liquid interface for at least 28 days before mucus collection. Moreover, as compared to primary cell-derived HAE, this provides a model which we could readily modify secreted mucin expression using CRISPR-Cas9. To evaluate the impact of airway mucins on NETosis, primary human neutrophils were isolated from whole blood samples from healthy human donors. As a renewable and readily available neutrophil model, we also used the human promyelocytic leukemia cell line HL-60 which can be differentiated (dHL-60) into a neutrophil-like state via exposure to low levels of dimethylsulfoxide (DMSO).^27^ dHL-60 cells have been used to study NETosis *in vitro* previously and can be evaluated using the same markers as PMNs.^28^ The wild type (WT) BCi-NS1.1 mucus was added to the media of polymorphonuclear leukocytes (PMNs) or dHL-60 cultures at doses normalized by the total protein concentration, up to 100 µg/mL. PMNs or dHL-60 cells were either stimulated using phorbol 12-myristate 13-acetate (PMA) to undergo NETosis or left unstimulated and treated with a vehicle to evaluate spontaneous NETosis. PMA is highly specific to the induction of classical lytic NETosis and does not have any known effects on other cell death pathways such as apoptosis in neutrophils.^24^

Sytox green is a membrane impermeable fluorescent dye that is frequently used to assess the extent of NETosis based upon staining of extracellular DNA. We found treatment with WT airway mucus significantly inhibits PMA-induced NETosis in dHL-60 cells based on Sytox green staining in fluorescent micrographs (**Figure 1A**). However, it should be noted mucus may also contain extracellular DNA that creates background fluorescence. Therefore, we used a microplate-based assay to quantify the fluorescence of Sytox green and background fluorescence originating from mucus-associated DNA was subtracted. This is similar to previously published studies and protocol articles employing Sytox-based microplate assays to assess NETosis.^29–32^ We treated dHL-60 cells with increasing concentrations of total mucus protein added to the media and found that WT airway mucus treatment inhibited PMA-induced NETosis based on the reduction in Sytox green fluorescence in a dose dependent manner (**Figure 1B**). This trend was mirrored in the unstimulated dHL-60 cells but was not statistically significant. To confirm these findings in native human neutrophils, primary human PMNs were isolated from whole blood using a density-gradient based separation method. The leukocytes were separated into two fractions, PMNs and peripheral blood mononuclear cells (PBMCs). To initially confirm that the isolated PMN samples were primarily granulocytes and PBMC samples were primarily monocytes and lymphocytes, we compared the CD14 expression (**Figure S1**). In the PBMC fraction, monocytes are typically CD14^high^, while other lymphocytes are CD14^-^. Granulocytes typically exhibit CD14^low^ expression.^33^ Neutrophils can be distinguished from other granulocytes if double positive for CD15 and CD16.^34^ We found that between 96-98% of the granulocyte population in the isolated PMN samples were CD15^+^CD16^+^ neutrophils (**Figure 1C**).

**Figure 1.**
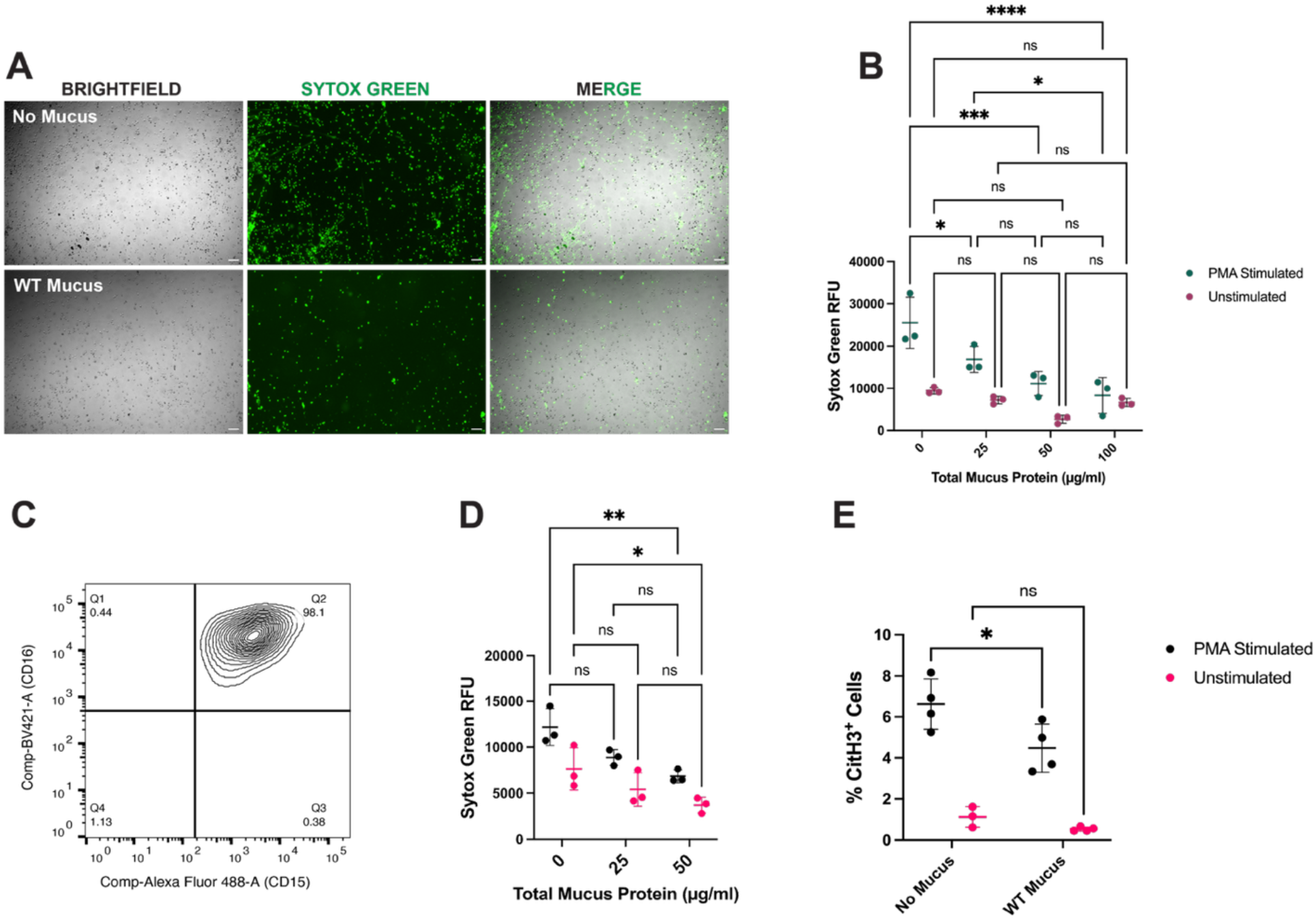
Airway mucus suppresses NETosis. A) Images of PMA-stimulated dHL-60 cells following 4 hours of incubation with no mucus or WT mucus (final concentration of 100 µg/ml total mucus protein) stained with Sytox green. Images captured at 5x magnification. Scale bars, 50 µm. B) Quantification of Sytox green fluorescence in relative fluorescence units (RFU) using a microplate assay. DHL-60 cells were treated with varying concentrations of WT mucus and either stimulated with PMA or left unstimulated. C) Flow cytometry plot showing the percentage of neutrophils (CD15^+^, CD16^+^) in the isolated PMN fraction from a healthy human blood donor. CD15 was detected using a primary antibody conjugated to Alexa Fluor 488 and CD16 was detected using a BV421-conjugated primary antibody. C) PMNs were treated with two different concentrations of total mucus protein and either stimulated with PMA to undergo NETosis or left unstimulated. Sytox green fluorescence of extracellular DNA was quantified using a microplate reader. D) CitH3 expression was quantified via flow cytometry. PMNs treated with 50 µg/ml total mucus protein were compared to those receiving no mucus treatment and were either stimulated with PMA or left unstimulated. * p < 0.05, ** p < 0.01, *** p < 0.001, **** p < 0.0001, ns = not significant. Statistical tests in B), D) and E) are two way ANOVAs with Šidák’s multiple comparisons test.

Following isolation, PMNs were treated with mucus to determine how it would affect NETosis. NETosis was evaluated using two different methods to quantify markers of NETs in PMNs. Quantification of Sytox green fluorescence revealed that extracellular DNA was significantly reduced in PMNs treated with increasing concentrations of WT mucus in both the PMA stimulated and unstimulated cells, signifying a reduction in both induced and spontaneous NETosis (**Figure 1D**). We also quantified the NETosis-specific protein marker citrullinated histone 3 (CitH3) via flow cytometry (**Figure 1E**). Representative flow cytometry plots for PMA stimulated PMN samples with versus without mucus treatment and the gating strategy and are shown in **Figure S2**. In agreement with the extracellular DNA results, we found that CitH3 expression was significantly reduced in PMA stimulated PNMs treated with 50 µg/ml WT mucus. This trend was also observed in the unstimulated PMNs, but was not significant. We compared this to CitH3 expression in the isolated PBMC fraction, since macrophages can also release extracellular traps.^35^ There were no significant effects of WT mucus on CitH3 in the PBMC fraction, suggesting that the effect of mucus is specific to NETosis (**Figure S3**). Primary HAE cells isolated from human donors are considered the gold standard of airway modeling *in vitro*. However as noted, we opted to use the immortalized BCI-NS1.1 cell line in this study to determine how genetic knockout of specific secreted mucins contributes to the inhibition of NETosis in subsequent experiments. However to confirm these findings with primary cell-derived HAE mucus, we compared the inhibition of NETosis in dHL-60 cells by WT BCI-NS1.1 versus WT mucus derived from primary HAE cells from a healthy donor to ensure similar effects. There was no significant difference in Sytox green fluorescence between mucus types (**Figure S4**).

### Loss of MUC5B or MUC5AC reduces inhibition of NETosis

Mucin glycoproteins were previously shown to be important in modulating NETosis in the oral cavity and female reproductive tract.^22–24^ Therefore, we characterized how the secreted airway mucin subtypes impacted NETosis inhibition. MUC5B and MUC5AC are the two major mucin subtypes that form the structure of the airway mucus gel.^1^ Previously published work from our lab used CRISPR-Cas9 to knockout (KO) expression of either MUC5B or MUC5AC in the BCI-NS1.1 HAE cell line.^36^ BCI-NS1.1 basal cells were previously transduced with a green fluorescent protein (GFP)-expressing lentiviral vector carrying both the single guide-RNA and Cas9 enzyme to knockout expression of either MUC5B (B KO) or MUC5AC (AC KO) and sorted using fluorescence activated cells sorting (FACS) for GFP^+^ cells.^36^ GFP^+^ cells were then cultured at air-liquid interface for at least 28 days until fully differentiated. Microscopy of the cell monolayer on day 30 of air-liquid interface culture confirmed GFP expression was retained in both the B KO and AC KO cultures (**Figure 2A**). In addition, we washed the cultures and collected the mucus to assess the relative amount of mucin in each sample using an assay to fluorometrically quantify O-linked glycosylation. We found that the WT mucus exhibited significantly higher fluorescence of O-linked glycans compared to the AC KO and B KO mucus samples (**Figure 2B**). The fold change in the fluorescence of the O-linked glycans was ∼0.7 in the AC KO and ∼0.6 in the B KO mucus compared to the WT. In addition, we wanted to specifically quantify the sialic acid glycans present in each sample. Sialic acid glycans were previously found to be primarily responsible for the ability of cervical mucins to inhibit NETosis.^22^ We found as compared to control (WT) mucus that there was a significantly lower concentration of sialic acid in the B KO mucus as compared to WT mucus (**Figure 2C**). A reduction in sialic acid was also apparent in the AC KO mucus as compared to WT mucus but these differences were not statistically significant.

**Figure 2.**
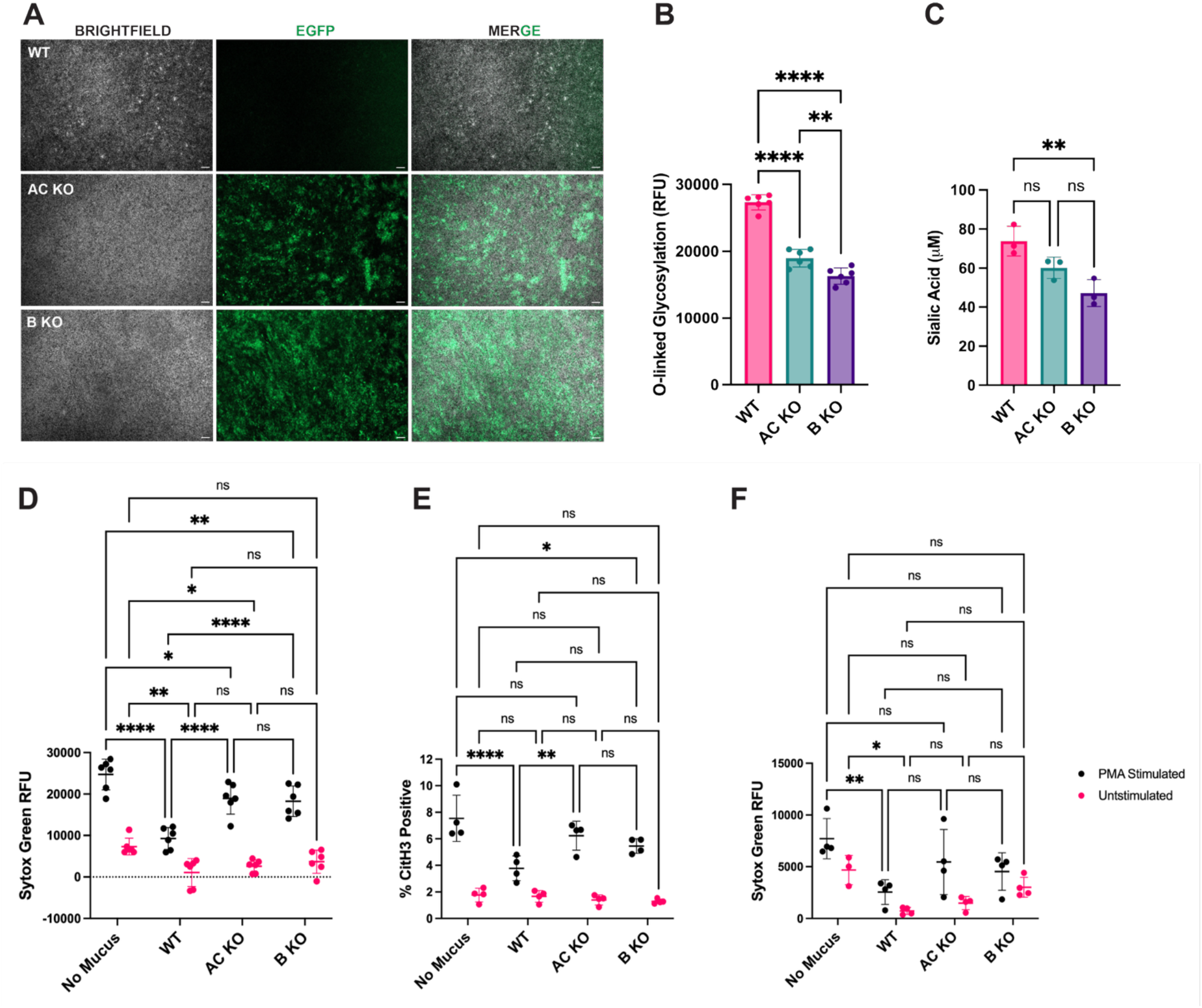
Mucus harvested from MUC5B and MUC5AC-deficient airway cultures has limited capacity to inhibit NETosis. A) Images comparing the GFP fluorescence of WT, B KO, and AC KO HAE cultures on day 30 of air-liquid interface culture at 5x magnification to demonstrate retention of the AC and B KO phenotypes. Scale bars, 50 µm. B) Fluorometric quantification of O-linked glycosylation in mucus samples from each HAE culture type. C) The concentration of sialic acid in WT, AC KO, or B KO mucus samples was quantified using a commercial kit. All mucus samples were diluted to 100 µg/ml total mucus protein. D) Sytox green fluorescence of dHL-60 cells treated with no mucus, or WT, AC KO, or B KO mucus samples at a final concentration of 25 µg/ml both with and without PMA stimulation. E) CitH3 flow cytometry of dHL-60 cells with and without PMA stimulation exposed to WT, AC KO, or B KO HAE mucus samples diluted to 25 µg/ml total mucus protein or no mucus treatment. F) Sytox green fluorescence of primary human PMNs treated with WT, AC KO, or B KO mucus (final concentration of 50 µg/ml total mucus protein) and stimulated with PMA or left unstimulated. * p < 0.05, ** p < 0.01, **** p < 0.0001, ns = not significant. Statistical comparisons in B) and C) are one way ANOVAs with Tukey’s multiple comparisons tests. Statistical tests performed in D), E) and F) are two way ANOVAs with Šidák’s multiple comparisons test.

We used the dHL-60 cell line to compare the effects of WT, B KO and AC KO mucus collected from BCI-NS1.1 cells on NETosis. In PMA-treated dHL-60 cells, both extracellular DNA and CitH3 markers of NETosis were reduced significantly by treatment with WT mucus in comparison to the B KO and AC KO mucus (**Figure 2D, E**). We confirmed that CitH3 was able to be detected in dHL-60 cells undergoing NETosis using the same staining protocol as PMNs (**Figure S5**). We quantified the fold change of both Sytox green fluorescence and the percent CitH3^+^ events for each experimental group receiving WT, AC KO or B KO mucus treatment compared to the no mucus treatment group. The fold changes in both markers were very similar between experiments in the cells induced with PMA (**Supplemental Table 1**). Differences in NETosis after treatment with the various mucus types were not as apparent in the unstimulated dHL-60 cells, likely due to the very low levels of spontaneous NETosis occurring. The effects of WT, B KO, and AC KO mucus on NETosis in PMNs follows a similar trend to the PMA-stimulated dHL-60 cells, in which WT was the only mucus type to produce a significant reduction in Sytox green fluorescence in both the PMA stimulated and unstimulated cells (**Figure 2F**).

In previously noted prior work, sialic acid glycans were found to be essential in the ability of purified bovine cervical mucins to suppress NETosis of bovine neutrophils.^22^ Salivary mucins are rich in Sialyl-Lewis X glycans that cause saliva to induce NETosis in human neutrophils.^23^ Both prior studies incorporated experiments employing a sialidase enzyme to cleave these key glycans and assess the impacts on NETosis. In addition, hypo-sialylation of mucus is characteristic of many chronic disease states and certain respiratory viruses like influenza.^37,38^ Considering these past studies, we de-sialylated mucus to give further mechanistic insight its ability to inhibit NETosis and explore how disease-associated alterations to mucus glycosylation may cause enhanced NETosis. Sialic acid concentration was depleted ∼4 fold in the sialidase-treated WT mucus compared to the mock-treated WT mucus (**Figure S6A**). The sialidase was heat inactivated before adding the de-sialylated mucus to dHL-60 cells to ensure that it did not interfere with the cellular glycocalyx. We also used a de-salting column to remove the cleaved sialic acid. The mock-treated mucus remained inhibitory of NETosis and significantly reduced Sytox green fluorescence in both the PMA stimulated and unstimulated dHL-60 cells compared to the no mucus control groups (**Figure S6B**). The cells exposed to sialidase-treated mucus exhibited increased fluorescence that was nearly identical to the no mucus control group, indicating loss of the ability of mucus to suppress NETosis without sialic acid. When we compared this to sialidase-treated mucus in which the cleaved glycans were not removed, both the sialidase and mock-treated mucus significantly reduced extracellular DNA fluorescence to the same extent compared to the no mucus control in the PMA stimulated groups (**Figure S6C**).

### NET formation is inhibited on mucus-secreting airway epithelium in a co-culture model

We wanted to confirm the importance of airway mucins in regulating NETosis when neutrophils are exposed to the natively secreted mucus gel on the epithelial surface. To accomplish this, we created a co-culture model in which dHL-60 cells were added directly to the mucosal surface of fully differentiated WT, AC KO, or B KO HAE cultures. After seeding the dHL-60 cells, we stimulated NETosis with PMA or left cells unstimulated and added a vehicle treatment instead. We stained cells with Sytox red, which was used instead of Sytox green to avoid interference with the GFP fluorescence of the underlying CRISPR-Cas9 edited cells in the AC KO and B KO cultures. We captured and analyzed images to quantify the mean fluorescence of the Sytox red after 4 hours of incubation in co-culture (**Figure 3A**). There was some background fluorescence from HAE culture-associated DNA, which was accounted for by quantifying the mean Sytox red fluorescence values obtained from images of WT, AC KO, or B KO cultures without dHL-60 cells. In agreement with our prior results with PMN/dHL60 stimulated with mucus added in a soluble form to the culture medium, we found significantly higher mean fluorescence of Sytox red when dHL-60 cells were seeded on either the AC KO and B KO HAE cultures compared to the WT (**Figure 3B, C**). This pattern occurred in both the PMA stimulated (**Figure 3B**) and unstimulated groups (**Figure 3C**), indicating enhanced induced and spontaneous NETosis when HAE cultures were deficient in either MUC5B or MUC5AC. In comparison, the HAE control monocultures showed similar mean Sytox red fluorescence intensities, with the AC KO and B KO having slightly lower fluorescence (**Figure 3D**). This result also confirmed that differences in the fluorescence were primarily due to changes in NETosis of dHL-60 cells and not DNA originating from HAE cultures.

**Figure 3.**
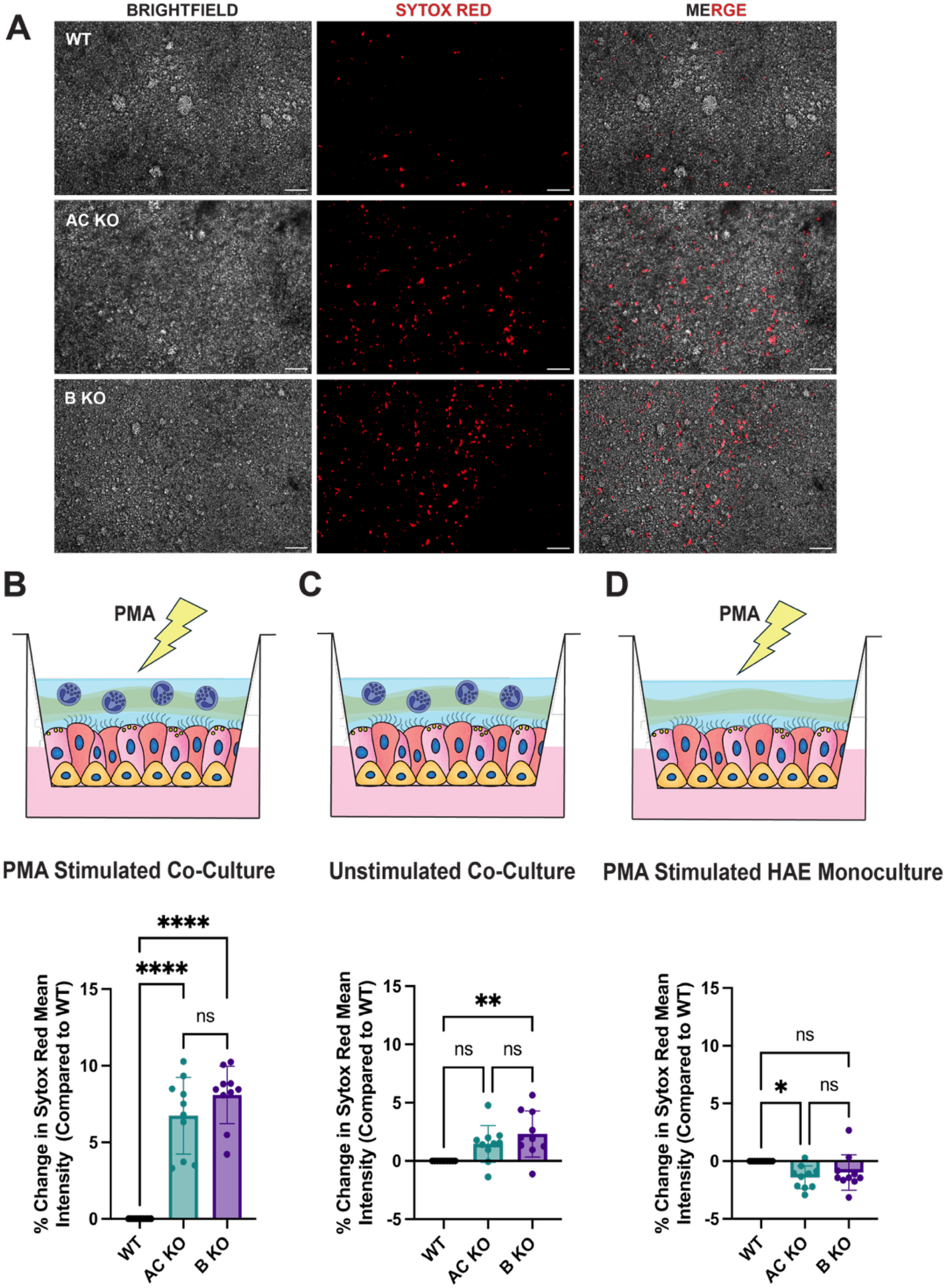
NET formation in the WT, MUC5B KO, and MUC5AC KO airway epithelial cell – HL60 co-culture model. A) Representative images of PMA stimulated dHL-60 cells in live co-culture with WT, AC KO, or B KO HAE cells following 4 hours of incubation taken at 10x magnification. Scale bars, 50 µm. B) Schematic of each co-culture or monoculture condition paired with the corresponding data quantifying the change in mean fluorescence intensity of Sytox red plotted directly below. * p < 0.05, ** p < 0.01, **** p < 0.0001, ns = not significant. Statistical tests performed in B), C) and D) are one way ANOVAs with Tukey’s multiple comparisons test post hoc.

### Stiffness-dependent regulation of NETosis in mucin-containing hydrogels

A prominent change to the mucus gel in many chronic pulmonary disease states is an increase in viscoelasticity. For example, mucus produced in the CF lung is hyper-concentrated leading to up to ∼100-fold increases in airway mucus viscoelasticity.^39^ In previous studies, NETosis was shown to increase as the stiffness of either polydimethylsiloxane (PDMS) or polyacrylamide-based hydrogel substrates increased.^40,41^ Considering this evidence in the context of muco-obstructive lung diseases, we investigated the effects of disease-associated mucus stiffening to determine how this may impact the inhibitory function against NETosis. To do this, hydrogel models were formulated to represent mucus secretions in healthy and diseased lungs. Our gel formulation approach was adapted from prior work from our lab by combining mucins with a thiolated 4-arm polyethylene glycol (PEG-4SH) to form disulfide bonds, as most reconstituted mucins will not undergo gelation alone.^42^ Prior to formulating the gels for these studies, we tested the effects of two different mucin types extracted from different animal tissues to ensure the mucins would produce the same inhibitory effect on NETosis. We compared porcine gastric mucins (PGM) to bovine submaxillary gland mucins (BSM) and found that PGM significantly reduced NETosis (**Figure S7**). BSM did not have significant effects on NETosis, which is in line since salivary mucins from the human oral cavity were previously shown to induce NETosis.^23^

We tested two PEG-4SH based hydrogel formulations in which we kept the concentration of PGM constant (0.5 mg/ml final concentration) and varied the concentration of the PEG-4SH crosslinker to be a final concentration of 35 mg/ml (3.5% PEG + PGM) or 50 mg/ml (5% PEG + PGM) to alter the gel stiffness (**Figure 4A**). We characterized the viscoelastic properties of the gels using particle tracking microrheology (PTM). Using this technique, the measured mean squared displacement (MSD) of nanoparticle probes diffusing in mucus and other soft biological materials can be used to characterize rheological properties as previously described.^21^ The 3.5% PEG + PGM gel demonstrated gel microrheological properties very similar to mucus isolated from primary HAE cultures from a healthy human donor (**Figure 4B-D**). By comparison, the 5% PEG + PGM gels exhibited a much denser microstructure with microrheological properties similar to what has been previously characterized in sputum samples of individuals with cystic fibrosis.^21^ The calculated pore sizes in the 5% PEG + PGM gel in particular is highly similar to what has been reported previously in cystic fibrosis sputum, which had an average pore size of 145 nm and a range of 60-300 nm.^43^

**Figure 4.**
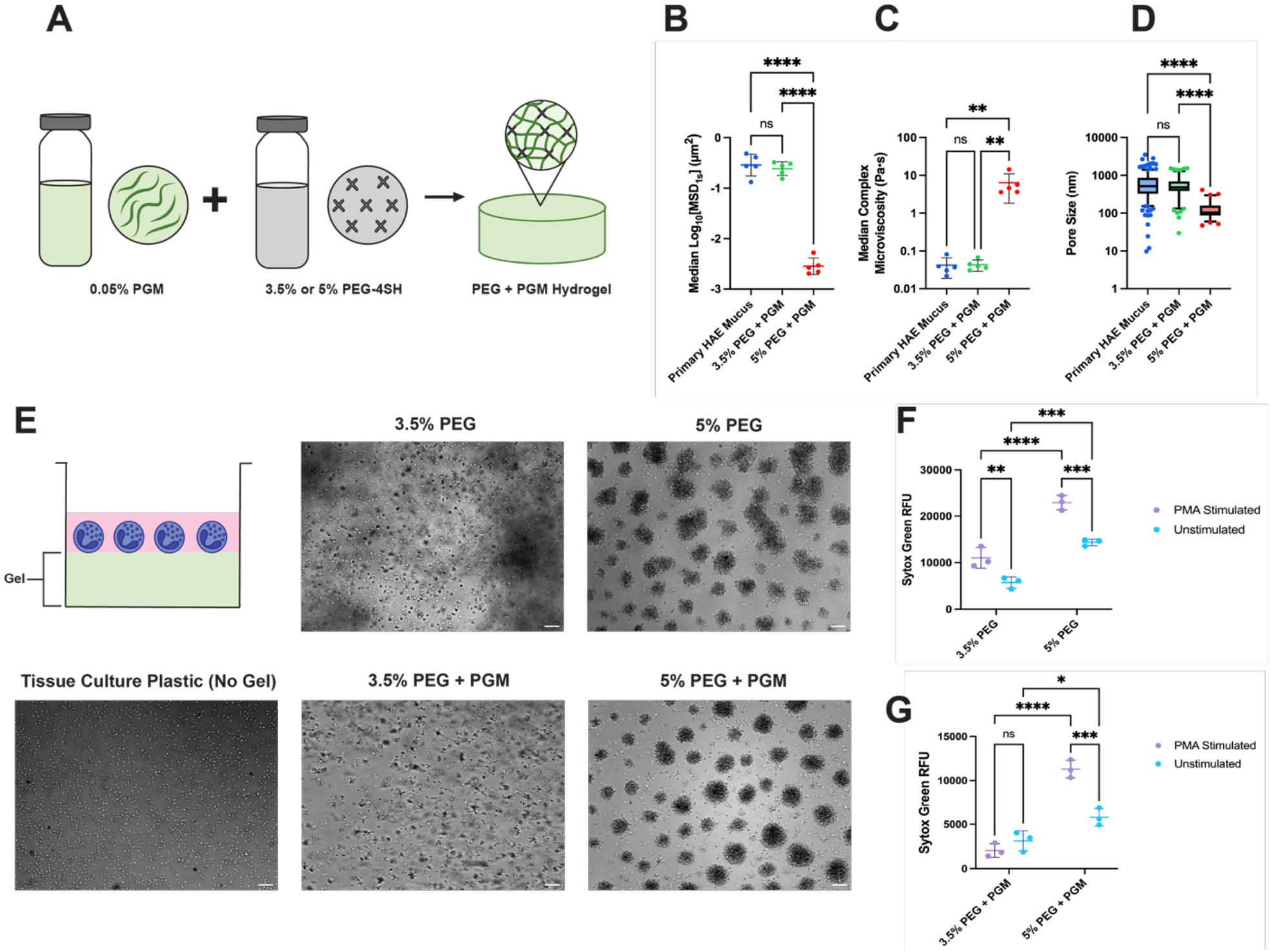
Mucin-containing hydrogel model with varied stiffness differentially regulates NETosis. A) Schematic figure representative of the PEG + PGM hydrogel formulation. Multiple particle tracking was used to quantify the microrheological properties of the 3.5% and 5% PEG + PGM gels compared to mucus collected from primary HAE cells from a healthy donor. B) The median values of the log_10_ of the particle MSD. The MSD values were used to calculate C) the complex microviscosity and D) pore sizes of each gel type or mucus sample. E) Schematic of the seeding of dHL-60 cells on gels and representative images of unstimulated dHL-60 distribution on the surface of each gel or tissue culture plastic following incubation for 4 hours imaged at 10x magnification. Scale bars, 50 µm. To evaluate the effect of gel stiffness on NETosis, the Sytox green fluorescence of dHL-60 cells seeded on each gel type was compared with and without PMA stimulation. F) The Sytox green fluorescence of dHL-60 cells seeded on 3.5% or 5% PEG gels with or without PMA stimulation. G) The Sytox green fluorescence of dHL-60 cells seeded on 3.5% PEG + PGM or 5% PEG + PGM gels with or without PMA stimulation. ** p < 0.01, *** p < 0.001, **** p < 0.0001, ns = not significant. Statistical tests performed in B), C) and D) are one way ANOVAs with Tukey’s multiple comparisons test and in F) and G) are two way ANOVAs with an uncorrected Fisher’s least significant difference post hoc test.

To determine the specific effects of mucins, we compared the 3.5% PEG + PGM and 5% PEG + PGM gels to those containing only PEG (3.5% PEG or 5% PEG) but otherwise formulated in the exact same manner. We seeded dHL-60 cells on the surface of each gel type and interestingly, we found that cells formed into clusters on the stiffer gels containing 5% PEG compared to the gels containing 3.5% PEG (**Figure 4E**). This effect was also observed in 5% PEG gels containing PGM. In the softer gels containing 3.5% PEG, the dHL-60 cells were able to penetrate and diffuse within the gels. The aggregation patterns were similar to what has been reported previously by monocytes seeded on collagen gels of varying stiffness, and indicate that this self-organization of dHL-60 cells is stiffness dependent.^44^

In order to determine if NETosis was affected by the difference in stiffness, we compared the Sytox green fluorescence of cells seeded on gels stimulated to undergo NETosis with PMA and unstimulated cells. In the gel formulations without PGM (3.5% PEG or 5% PEG), the fluorescence was significantly increased with PMA stimulation in both formulations, with a ∼1.9 fold increase in fluorescence values with PMA stimulation compared to unstimulated cells seeded on the 3.5% PEG gels and a ∼1.6 fold increase in fluorescence values with PMA stimulation compared to unstimulated cells seeded on the 5% PEG gels (**Figure 4F**). Overall, we found that PMA treatment resulted in significant upregulation of NETosis in both the soft (3.5%) and stiff (5%) PEG gels without PGM added. In addition, there was a ∼2.1 fold increase in Sytox green fluorescence in the PMA-stimulated cells seeded on the 5% PEG gel compared to the 3.5% PEG gel and a ∼2.5 fold increase in the fluorescence of the unstimulated cells seeded on the 5% PEG gels compared to the 3.5% PEG gels. This indicates that increased PEG gel stiffness resulted in more NETosis, which is in agreement with previous studies of the effects of gel stiffness on NETosis.^40,41^ However, there is a potential caveat in comparing NETosis on the soft and stiff gel formulations as the cells’ are more able to migrate within the soft gels and generally remain on the gel surface in the stiff gel may affect the fluorescence. In addition, stiffer gels often induce more apoptosis in leukocytes such as monocytes.^44^

We also compared this to the induction of NETosis when dHL-60 cells were seeded on gels containing PGM (**Figure 4G**). When cells were seeded on the softer 3.5% PEG + PGM gel, there was no change in Sytox green fluorescence between the PMA stimulated cells and unstimulated cells. This indicated that NETosis was successfully inhibited in the PMA stimulated cells, likely because the cells were able to migrate within the softer gel and interact with mucins. However, when cells were seeded on the stiffer 5% PEG + PGM gel, Sytox green fluorescence increased ∼1.9 fold compared to the unstimulated cells. NETosis was not inhibited significantly when cells were seeded on the stiffer 5% PEG + PGM gel, potentially due to the inability of the cells to interact with mucins incorporated into the gel. Similar to the PEG-only gels, we also found a greater extent of NETosis for dHL-60 when cultured on the stiffer 5% PEG + PGM gel compared to the softer 3.5% PEG + PGM hydrogels. The Sytox green fluorescence was significantly increased for dHL-60 cells cultured on the 5% PGM + PEG gels compared to the 3.5% PEG + PGM gels in both the PMA stimulated and unstimulated groups.

## Discussion

Neutrophils recruited to mucosal tissues must balance inflammatory responses against infection with regulatory checkpoints to limit uncontrolled inflammation and excessive tissue damage. It has been shown previously that mucin glycoproteins from the oral cavity and female reproductive mucosal tissues are highly immunomodulatory and can regulate NETosis.^22–24^ However, the mucins from each mucosal tissue type have been found to differentially regulate NETosis to help maintain neutrophil homeostasis where tissue-specific mucin-associated glycan profiles have a well-documented role in directing neutrophil fate. For example, it has been shown that saliva in the oral cavity induces NETosis via interactions with the sialyl-Lewis X glycans of mucins. One could reason the function of saliva in activating NETosis is due to the extremely high density of microbes present in the mouth. A reduced capacity of saliva to induce NETosis is also associated with the development of oral ulcers.^23^ In contrast, another study found that mucins purified from bovine cervices inhibited NETosis via sialic acid glycans. Increased NETosis is also associated with various cervical and pregnancy related disease-states.^45^ In this work, we determined the role of airway mucus and secreted mucin subtypes in maintaining neutrophil homeostasis in the airways. In addition, we investigated how alterations to the physicochemical properties of mucus may lead to dysregulated NETosis.

We found that WT HAE mucus was able to strongly inhibit NETosis in both primary human PMNs and in the neutrophil model cell line dHL-60. This aligns with what is observed clinically as NETs are rarely observed in the lumen of healthy airways, but enhanced NETosis is often characteristic of both chronic lung diseases and acute respiratory infection in which the mucus composition is altered.^6,10–13^ To determine the role of airway-associated MUC5B and MUC5AC mucin subtypes, we used HAE cells genetically engineered to knockout the expression of either mucin. We found that loss of either MUC5B or MUC5AC enhanced NETosis compared to the WT mucus. We also wanted to ensure that the role of secreted MUC5B and MUC5AC mucins remained important in regulating NETosis in a more complex model incorporating the epithelium. Therefore, we demonstrated that the regulatory function of airway mucins in NETosis is maintained for neutrophils in contact with a mucus-coated airway epithelium *in vitro* using a co-culture model. These effects were diminished in the co-culture model in HAE cells that lacked expression of either MUC5B or MUC5AC, showing each airway mucin may serve as a regulator of NET formation.

Given the similarities between airway mucins and cervical mucins in reducing NETosis, we investigated the role of sialic acid glycans on NET inhibition by removing these functional groups via neuraminidase treatment. Our data suggest that sialic acid terminated glycans on airway mucin glycoproteins interact with sialic acid binding immunoglobulin-like lectins (siglecs) expressed on the surface of neutrophils. In particular, siglec 9 is highly expressed by human neutrophils and has immune receptor tyrosine-based inhibitory motifs (ITIM) domains.^46^ Engagement of siglec 9 has previously been shown to be inhibitory of NETosis.^28,47^ Airway mucins have been identified as siglec 9 ligands in prior literature, making it highly likely that the interaction between siglec 9 and mucin-associated sialic acid is the specific pathway by which NETosis is reduced by airway mucus.^48^ Siglec 9 expression on neutrophils is also upregulated in individuals with acute COPD exacerbations and given the elevated concentrations of NETs in the COPD lung, upregulation of siglec 9 expression may provide a means to counteract excessive NETosis.^49^

The increase in NETosis that we observed when mucus was treated with sialidase also gives insight into how aberrant mucus sialylation in chronic and acute respiratory disease may contribute to enhanced NETosis. For example, hypo-sialylation of mucins is observed in cystic fibrosis due to reduced sialyltransferase expression.^37^ Low charge mucin glycoforms are observed in several other chronic lung disease states including COPD and asthma.^50,51^ In addition, respiratory pathogens can alter glycosylation of mucins in the airway. Influenza virus expresses neuraminidase that cleaves sialic acid glycans, and infection is associated with increased pulmonary NETosis *in vivo*.^15^ It was previously found that the metabolite pyocyanin, secreted by *Pseudomonas aeruginosa*, significantly increased sialyl-Lewis X glycosylation of MUC5AC secreted by HAE cells.^52^ If the increase is very substantial, it is possible that the upregulation of sialyl-Lewis X glycosylation may contribute to enhanced NETosis in *Pseudomonas aeruginosa* infections of the lungs given its role in inducing NETosis in saliva.

Another important alteration in airway secretions observed in chronic lung diseases is the build-up of highly concentrated mucus that possesses significant alterations in viscoelasticity. As NETosis has previously been shown to be a mechano-sensitive process, we used a hydrogel model to evaluate stiffness-dependent activation of NETosis. While the dHL-60 cells did not undergo increased NETosis with PMA stimulation compared to the unstimulated cells in the representative healthy-state gels (3.5% PEG + PGM), there was a significant increase in NETosis when dHL-60 cells seeded on the representative disease-state gels (5% PEG + PGM) were stimulated with PMA compared to the unstimulated cells. In contrast, when mucin was removed and gels consisted only of PEG-4SH, dHL-60 NETosis was significantly increased with PMA stimulation compared to the unstimulated cells on both the softer 3.5% PEG and stiffer 5% PEG gels. We interpret these results as evidence that migration of dHL-60 cells within the gels is required for the neutrophils to engage receptors on mucin glycoproteins to modulate NETosis. In the context of the *in vivo* environment, NETs are often prominent in highly viscoelastic mucus plugs that form in chronic lung diseases. In one study where they evaluated mucus plugs in asthma and COPD, there were significant increases in DNA and CitH3 expression, indicating more NETosis.^53^ Our experiments using the healthy and disease-like hydrogel formulations were performed as proof-of-concept that disease-associated increases in stiffness may interfere with mucin-dependent inhibition of NETosis in the airways. Future studies investigating this concept in native mucus with varied stiffness will be performed to determine if this is indeed a mechanism by which NETosis is upregulated in disease.

In summary, we have identified secreted mucus in the airway as an endogenous regulator of NETosis. Airway mucus inhibits NETosis in a sialic acid-dependent manner, aligning with other studies demonstrating the importance of mucin-associated glycans in modulating NETosis in mucosal tissues. Our data also suggest that the changes to the biophysical properties of mucus in airway diseases such as asthma, CF, and COPD may impede inhibition of NETosis. Future studies will further investigate both aberrant glycosylation and biophysical properties as potential mechanisms by which the mucus loses its immunomodulatory ability. Our findings on mucin– regulated NETosis in health and disease may guide new therapeutic strategies to restore mucus homeostasis and prevent pathological NET release in respiratory diseases.

## Methods

### PMN and PBMC Isolation and Flow Cytometry

Whole blood samples from screened healthy human donors were obtained from the American Red Cross. Blood components were separated using a density gradient based approach with Polymorphprep reagent based on the manufacturer’s instructions (Progen). 6 ml of Polymorphprep was added to a 15 ml conical tube, then 6 ml of human whole blood was carefully layered on top. The tubes containing the Polymorph prep and blood were centrifuged at room temperature at 500xg for 30 minutes to separate the blood components. The centrifuge deceleration brake was disabled to prevent disruption of the density gradient. The density gradients were carefully unloaded and both the PMN and PBMC fractions were collected. The PMNs and PBMCs were washed in sterile PBS. After washing, PBMCs were resuspended in serum free RPMI 1640 media. PMNs were resuspended in 2 ml of red blood cell lysis buffer (eBioscience 1X RBC Lysis Buffer) for 30 seconds, then diluted in 10 ml serum free RPMI 1640 media. PMNs were centrifuged and red blood cell lysis was repeated once more if there was visible red blood cell contamination of the pellet. PMNs were then resuspended in serum free RPMI media. Both the PBMCs and PMNs were counted and diluted to the desired cell concentration.

To evaluate the success of the PMN isolation, we used flow cytometry to check for the surface markers CD14, CD15, and CD16. We compared the presence of these markers to the PBMC fraction. To stain the PMNs and PBMCs, 500 µL of either cell suspension diluted to 10^6^ cells/ml (5×10^5^ cells total) in serum free RPMI 1640 media was aliquoted into microcentrifuge tubes. Cells were blocked in 100 µL of human TruStain FcX Fc Receptor Blocking reagent (BioLegend) diluted 1:100 in a solution of 2% heat inactivated fetal bovine serum in sterile PBS (FACS buffer). Both PMNs and PBMCs were blocked in the Trustain and FACS buffer solution for 15 minutes on ice. Following blocking, cells were pelleted and resuspended in 100 µL of PBS. To stain for CD14, CD15, and CD16 three primary antibodies conjugated to different fluorophores were used (BioLegend 301813 (CD14), BioLegend 301910 (CD15), BioLegend 302037 (CD16)). 2.5 µL of each antibody was added to the cell suspension in PBS. Both single antibody-stained controls and unstained controls were prepared in parallel. Cells incubated with surface marker antibodies for 30 minutes on ice protected from light. Cells were washed in PBS and fixed for 25 minutes in 2% paraformaldehyde at 4°C protected from light. Following fixation cells were washed again in PBS and resuspended in 400 µL of PBS. Flow cytometry was performed on the stained PMN and PBMC samples. Populations were gated and graphs were created using FlowJo (version 10) software.

### HL-60 Culture and Differentiation

The immortalized human acute myeloid leukemia cell line HL-60 (ATCC CCL-240) was cultured in a flask in RPMI 1640 media supplemented with 20% heat inactivated fetal bovine serum and 1% penicillin-streptomycin (complete RPMI 1640 media).^28^ The HL-60 cells were cultured for 2 weeks prior to beginning differentiation into a neutrophil-like state and media was replaced every 2-3 days. To begin differentiation into dHL-60 cells, 1.3% DMSO was added to the complete RPMI 1640 media for a period of 5-7 days.^28,54^ The DMSO-containing complete RPMI 1640 media with was refreshed 48 hours prior to using cells for experiments.

### WT and KO HAE Cell Culture

The immortalized human airway epithelial basal cell line BCI-NS1.1 (gifted by Ronald Crystal, Cornell University) were grown in a flask with Pneumacult Ex-Plus media (Stemcell Technologies) until 70-80% confluent. To begin basal cell differentiation, cells were seeded at a density of 10^4^ cells/cm^2^ on collagen-coated Transwell membranes in a 12 well plate (Corning, 6.5 mm). Pneumacult Ex-Plus media was added to apical and basolateral compartments of the Transwell inserts and cells were kept in submerged culture until reaching confluency. Once a monolayer of basal cells formed on the Transwell membrane, cells were airlifted by removing the media from the apical compartment to establish air-liquid interface (ALI) cultures. The media in the basolateral compartment was switched to Pneumacult ALI (Stemcell Technologies) when cells were airlifted. Media was replaced in the basolateral compartment every other day and cells were cultured at ALI for 28 days until fully differentiated into a mucociliated epithelium.

The MUC5B KO (B KO) and MUC5AC KO (AC KO) BCI-NS1.1 cell lines were previously created and characterized by our lab.^36^ For each mucin type, two different single guide RNAs were created to target two different exon regions of MUC5B or MUC5AC, termed MUC5B/AC KO1 or KO2. In this work, we used the MUC5AC KO1 and MUC5B KO2 lines, with the specific guide RNA sequences previously published.^36^ After thawing the cryopreserved cell lines, we expanded the basal cells in a flask. We sorted the cells for GFP expression using FACS to ensure a pure population and cultured the sorted cells in a flask. The AC KO and B KO cells were seeded on Transwell inserts, airlifted, and cultured at ALI using the same methods described above.

### HAE Mucus Collection and Quantification of Total Protein

Mucus was washed and collected from the fully differentiated WT, B KO, and AC KO HAE cultures after 28 days at ALI. The WT, B KO, and AC KO HAE cells were all kept on the same washing schedule prior to ensure similar mucus secretion patterns. To remove mucus from the surface of the HAE cultures, 250 µL of PBS was added to the apical compartment of HAE cultures and placed back into the incubator (37°C, 5% CO_2_) for 30 minutes. The washings were collected from the surface of the cultures using a pipette and pooled together. 500 µL of the collected washings was added to 0.5 ml Amicon Ultra 100 kDa filters (Millipore Sigma) and centrifuged at 14,000xg for 20 minutes to remove the PBS. The mucus was stored at -80°C immediately following centrifugation and thawed at 4°C for use in experiments. If larger volumes of mucus were necessary for an experiment, multiple mucus samples from WT, B KO or AC KO HAE cultures were thawed and pooled together. A bicinchonic acid (BCA) assay was performed on the pooled mucus to determine the total protein content of each sample and inform the mucus dilution factors to ensure the same total mucus protein concentration in subsequent experiments. The BCA assay (Thermo Fisher Scientific Pierce BCA) was performed according to kit instructions.

### Microplate Assay to Evaluate Extracellular DNA (PMN and dHL-60)

A BCA assay was performed to determine total mucus protein concentration of each sample prior to beginning the microplate assay. dHL-60 cells or primary PMNs were harvested and resuspended in serum free RPMI 1640 media. Based on the BCA assay results, the appropriate volume of each mucus sample type was added to aliquots of PMNs or dHL-60 cells diluted in serum free RPMI 1640 media to achieve final concentrations of 10^6^ cells/ml and 50 µg/ml total mucus protein for PMN experiments or 10^6^ cells/ml and 25 µg/ml total mucus protein for dHL-60 experiments. In control groups that did not receive any mucus treatment PMNs or dHL-60 cells were diluted to 10^6^ cells/ml in serum free RPMI media. Aliquots containing only mucus (no cells) diluted to the same final concentrations in serum free RPMI 1640 media were prepared in parallel to control for background Sytox fluorescence from mucus-associated extracellular DNA. 100 µL of each aliquot was added to the wells of a black optical bottom 96 well plate (Thermo Scientific Nunc) to achieve a seeding density of 10^5^ cells/well. Cells were incubated for 30 minutes at 37°C and 5% CO_2_ prior to stimulation of NETosis with PMA or treatment with an equal volume of a vehicle in unstimulated groups. In PMN experiments, a final concentration of 50 nM PMA was added to each well and a final concentration of PMA was 100 nM was added in dHL-60 experiments. Cells were incubated for 4 hours at 37°C and 5% CO_2_. Sytox green (Invitrogen) was diluted to a concentration of 0.5 µM in the PMN experiments and 1 µM in dHL-60 experiments in sterile PBS. 10% v/v of the Sytox green solution was added to each well, to achieve a final concentration of 0.05 µM in PMN experiments and 0.1 in dHL-60 experiments. Cells were incubated for 15 minutes at 37°C and 5% CO_2_. The Sytox green fluorescence was read on a microplate reader at excitation and emission wavelengths of 504 nm and 523 nm, respectively.

### CitH3 Staining and Flow Cytometry (PMN and dHL-60)

A BCA assay was first conducted to determine the total mucus protein concentration of each sample and calculate dilution factors. PMNs or dHL-60 cells resuspended in serum free RPMI 1640 media were diluted to final concentrations of 10^6^ cells/ml and 50 µg/ml total mucus protein for PMN experiments or 10^6^ cells/ml and 25 µg/ml total mucus protein for dHL-60 experiments. In samples that did not receive mucus treatment, cells were diluted to 10^6^ cells/ml in serum free RPMI media. 1 ml of each cell suspension was added into the wells of a 12 well tissue culture plate (final seeding density of 10^6^ cells/well) and incubated for 30 minutes at 37°C and 5% CO_2_. PMA was added to each well at a final concentration of 50 nM for PMN experiments and 100 nM for dHL-60 experiments. In unstimulated wells, an equal volume of the vehicle was added instead of PMA. Cells were incubated for 4 hours at 37°C and 5% CO_2_ to allow NETosis to occur. The contents of each well were collected by pipetting up and down to dislodge cells from the bottom of each well, then transferring to microcentrifuge tubes. However, cells in advanced stages of NETosis appeared to be more adherent to the bottom of the wells when observed under the microscope and were not dislodged by pipetting alone. Therefore, after pipetting and collecting the initial cell suspension we added additional serum free RPMI 1640 media to the wells and used a cell scraper to remove any residual cells and NET debris from the wells. The additional media solution was then collected in the same microcentrifuge tube.

PMN or dHL-60 cells were fixed for 20 minutes in 4% PFA at 4°C, then washed in PBS. Cells were blocked in 100 µL of human TruStain FcX Fc Receptor Blocking reagent (BioLegend) diluted 1:100 in a solution of 2% heat inactivated fetal bovine serum in sterile PBS (FACS buffer) for 30 minutes at 4°C and washed. To stain cells for extracellular CitH3, we resuspended cells in 100 µL of a primary rabbit anti-human CitH3 antibody diluted 1:200 in PBS (Abcam 5103). Cells incubated with the primary antibody for 30 minutes at 4°C and were washed in PBS. Cells were resuspended in 100 µL of solution of a secondary donkey anti-rabbit antibody conjugated to Alexa Fluor 647 diluted 1:200 in PBS (BioLegend Poly4064). Both unstained controls and controls in which the cells were incubated with only the secondary antibody were prepared in parallel to account for any non-specific binding. Cells were incubated with the secondary antibody for 30 minutes protected from light at 4°C, then washed in PBS. Finally, cells were resuspended in 300 µL of PBS. Flow cytometry on each PMN or dHL-60 sample was performed, and populations were gated using FlowJo software (version 10).

### O-linked Glycosylation Fluorometric Assay

The relative mucin content of WT, B KO, and AC KO mucus samples was assessed using a previously established assay to fluorometrically quantify O-linked glycans.^55^ A BCA assay was performed prior and all mucus samples were diluted to a final concentration of 100 µg/ml total mucus protein in PBS. Solutions of 0.6 M 2-cyanoacetamide (CNA) and 0.15 M sodium hydroxide (NaOH) were combined to yield a solution with a final concentration of 0.1 M CNA and 0.025 M NaOH. 60µL of this CNA/NaOH reagent was added to 50 µL of each diluted mucus sample and incubated on a heat block at 100°C for 30 minutes. The reaction was quenched by adding 500 µL of a 0.6 M boric acid buffer (pH 8.0) to each sample. Samples were added to the wells of a back 96 well plate and the fluorescence was quantified using a microplate reader with excitation and emission wavelengths of 336 and 383 nm, respectively.

### HAE and dHL-60 Co-Culture and Image Analysis

dHL-60 cells were diluted to 1.5×10^6^ cells/ml in sterile PBS. Sytox red (Invitrogen) was added to the cell suspension to a final concentration of 0.05 µM. Blank PBS containing the same dilution of Sytox red without dHL-60 cells was also prepared to be applied to control cultures. 200 µL of the dHL-60 cell suspension or the blank PBS was added to the apical chamber of HAE cells (final seeding density of 3×10^5^ dHL-60 cells). The cultures were incubated for 30 minutes at 37°C and 5% CO_2_. PMA was added to the apical compartment to a final concentration of 100 nM based on the volume of the cell suspension or blank PBS added to induce NETosis. In unstimulated cultures, an equal volume of the vehicle was added. Cultures were incubated for 4 hours at 37°C and 5% CO_2_.

All HAE cells had been washed 1 week prior to allow for sufficient mucus accumulation before co-culture. In addition, it is impossible that the dHL-60 cells would have transmigrated across the airway epithelial layer, as the pore sizes of the Transwell membranes were 0.4 µm. The apical surface of each HAE culture was imaged at 10x magnification (Zeiss LM 800) using excitation and emission wavelengths of 640 and 658 nm, respectively, to detect Sytox red fluorescence. Images were analyzed using FIJI software. All images were set to the same minimum and maximum pixel intensity values, then a threshold was applied to exclude background fluorescence. The mean pixel intensity of the thresholded images was quantified and the percent change in the mean pixel intensity of the B KO and AC KO cultures compared to the WT was calculated. The control cultures without dHL-60 cells were also analyzed using the same thresholding method to compare residual background fluorescence in each culture type and the percent change in the mean pixel intensity of the B KO and AC KO cultures compared to the WT was calculated.

### Mucus De-Sialylation and Sialic Acid Quantification

A BCA assay was performed to determine total mucus protein concentration, then WT mucus was diluted to 250 µg/ml in a solution of 1 mM CaCl_2_. The sialidase enzyme used was neuraminidase purified from *Vibrio* cholerae (Sigma-Aldrich N7885) was diluted in a solution of 1 mM CaCl_2_ to a final concentration of 1000 mU/ml. For mock treated mucus, a vehicle was diluted in 1 mM CaCl_2_.The diluted mucus solution was combined with the sialidase enzyme solution for a final concentration of 100 mU/ml or an equal volume of the mock solution. The mock and sialidase treated mucus was incubated overnight at 37°C. To inactivate the neuraminidase, samples were placed on the heat block at 80°C for 30 minutes.^56^ After this, the cleaved glycans were removed and the CaCl_2_ buffer was exchanged using a 7 kDa molecular weight cut off Zeba de-salting column. The de-salting column was used according to manufacturer’s instructions, and samples were buffer exchanged into sterile PBS. Following sample preparation, mucus was diluted to a final total protein concentration of 25 µg/ml in subsequent dHL-60 microplate assays to quantify NETosis via Sytox green fluorescence.

Sialic acid concentration of the de-sialylated versus mock treated mucus and the WT, B KO and AC KO was determined using a commercial kit (Sigma-Aldrich MAK314). All samples were diluted to 100 µg/ml total mucus protein in PBS according to the results of a prior BCA assay. The manufacturer’s protocol was then followed to quantify the total sialic acid in each sample. Briefly, sialic acid is oxidized formylpyruvic acid that reacts with thiobarbituric acid to yield a fluorometric product. The fluorescence of each sample was quantified in a black 96 well plate using a microplate reader at an excitation and emission wavelength of 555 nm and 585 nm, respectively. A standard curve was also prepared to quantify the concentration of sialic acid in each sample.

### Synthetic Hydrogel Preparation and Extracellular DNA Assay

Porcine gastric mucins (PGM) (Sigma; Type III) were dissolved in PBS at a final concentration of 1 mg/ml and stirred for 1 hour. PEG-4SH (Laysan Bio; 10kDa) was dissolved separately in PBS at a concentration of either 70 mg/ml or 100 mg/ml. Both the mucin solution and each PEG-4SH solution were sterile filtered (0.45 µm pore size) in a biosafety cabinet to remove contaminants. The gels were prepared by combining equal volumes of the 1 mg/ml mucin and 70 mg/ml PEG-4SH or 100 mg/ml PEG-4SH. This yielded gels with final concentrations of 0.5 mg/ml PGM (0.05%) and 35 mg/ml PEG-4SH (3.5% PEG) to model healthy state mucus or 0.05% PGM and 50 mg/ml PEG-4SH (5% PEG) to model disease state mucus. 100 µL of each gel solution was added to the wells of a black optical bottom 96 well plate. The lid was placed back on the 96 well plate and sealed with parafilm. Gelation occurred over 48 hours at room temperature.

After gels were set, a similar procedure to what is described above was followed to detect NETosis using Sytox green fluorescence. dHL-60 cells were diluted to a final concentration of 10^6^ cells/ml in serum free RPMI 1640 media. 100 µL of the cell suspension was added to the wells containing the gels (final seeding density of 10^5^ cells/well). Background fluorescence was accounted for using control wells in which 100 µL of blank media without dHL-60 cells was added to the wells with various gel formulations. Plates incubated for 30 minutes at 37°C and 5% CO_2_ following cell seeding. NETosis was stimulated in certain wells using 100 nM final concentration of PMA, based on the volume of 100 µL of cell suspension added to each well. In unstimulated wells, cells were treated with a vehicle. Cells were stained with a final concentration of 0.05 µM in each well after 4 hours of incubation at 37°C and 5% CO_2_. The plates were incubated with the Sytox green stain for 15 minutes and the Sytox green fluorescence was read using a microplate reader with excitation and emission wavelengths of 504 nm and 523 nm, respectively.

### Multiple Particle Tracking Microrheology

Gel solutions were prepared in the same manner as described above and added to microscopy chambers. The microscopy chambers were created by adhering an O-ring to the surface of a glass microscope slide using vacuum grease, forming a shallow well for the gel to form. 25 µL of each gel solution was added to each microscopy chamber. Mucus collected from primary human airway epithelial cells was used as a comparison to determine the physiological relevance of the gel properties. The same procedures were used in preparing the mucus samples for multiple particle tracking as the gel solutions. Red fluorescent 100 nm polystyrene nanoparticles (Life Technologies Fluospheres) were densely coated in 5kDa PEG to minimize electrostatic interactions with the gel or mucus matrices as previously described.^42^ 1 µL of PEGylated nanoparticles diluted 1:500 in PBS was added to the gel solution. The microscope slides were placed inside a petri dish with a damp Kim-wipe to prevent the gels from drying out, and the dish was sealed with parafilm. Gelation with the nanoparticles added occurred over 24 hours at room temperature.

Multiple particle tracking and analysis was performed as previously described.^21,42,57^ Briefly, fluorescent PEGylated nanoparticles diffusing within the gels or mucus were imaged at 63x magnification (Zeiss LSM 800). Videos of the nanoparticles were taken in multiple regions of each gel or mucus sample. The videos were 10 seconds in length with a frame rate of 33 Hz. The material properties of the gels or mucus samples were determined based on the nanoparticle mean squared displacement (MSD). A MATLAB algorithm was used to identify and track the MSD of the nanoparticles in each video as a function of the lag time (τ) with the equation ⟨Δ*r*^2^(*τ*)⟩ = ⟨(*x*^2^ + *y*^2^)⟩. Previous work found that static and dynamic error in multiple particle tracking experiments was reduced at *τ* = 1*s* (MSD_1s_) and we therefore reported the median MSD_1s_ in this work.^58,59^

The MSD values of each particle could then be used to calculate the microrheological properties of the gel or mucus samples. The generalized Stokes Einstein relation is used to relate the viscoelastic spectrum [*G*(*s*)] to the Laplace transform of ⟨Δ*r*^2^(*τ*)⟩, which is ⟨Δ*r*^2^(*s*)⟩. The generalized Stokes Einstein relation is defined as *G*(*s*) = 2*k*_*B*_*T*/[*πa*⟨Δ*r*^2^(*s*)⟩] where *k*_*B*_*T* is thermal energy, *a* is the radius of the particles, and *s* is the complex Laplace frequency. The *s* variable can be substituted with *iω*, where *i* is a complex number and *ω* is frequency, to compute the complex modulus using the equation *G*^∗^(*ω*) = *G*^′^(*ω*) + *G*^′′^(*iω*). The complex microviscosity (*η*^∗^) can then be calculated using the complex modulus using the equation *η*^∗^(*ω*) = *G*^∗^(*ω*)/*ω* and considers both the elastic (*G*^′^) and viscous (*G*^′′^) moduli. Based on the values of the elastic modulus (*G*^′^) the pore size of the gel matrices can be calculated as 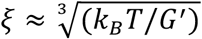.

## Acknowledgements

This project was funded by the Cystic Fibrosis Foundation (DUNCAN24G0) and National Institutes of Health (R01 HL160540, R01 HL151840, F31HL176146).

## Author contributions

A.B. and G.A.D. conceptualized the project and designed experiments. A.B and V.R. conducted experiments and led data analysis. A.B. and G.A.D. wrote the article. All authors have given approval to the final version of the manuscript.

## References

1. Song, D., Cahn, D. & Duncan, G. A. Mucin Biopolymers and Their Barrier Function at Airway Surfaces. Langmuir 36, 12773–12783 (2020).

2. Belda, J. et al. Induced sputum cell counts in healthy adults. Am J Respir Crit Care Med 161, 475–478 (2000).

3. Frey, D. L. et al. SputOMICs identifies common and distinct markers in cystic fibrosis and chronic obstructive pulmonary disease. Sci Rep 15, 44418 (2025).

4. John V. Fahya, B., Kimb, K. W., Liub, J. & Homer A. Bousheya, B. Prominent neutrophilic inflammation in sputum from subjects with asthma exacerbation. Journal of Allergy and Clinical Immunology 95, 843–852 (1995).

5. Johansson, C. & Kirsebom, F. C. M. Neutrophils in respiratory viral infections. Mucosal Immunol 14, 815–827 (2021).

6. Pan, T. & Lee, J. W. A crucial role of neutrophil extracellular traps in pulmonary infectious diseases. Chin Med J Pulm Crit Care Med 2, 34–41 (2024).

7. Rosales, C. Neutrophil: A Cell with Many Roles in Inflammation or Several Cell Types? Front Physiol 9, 113 (2018).

8. Brinkmann, V. et al. Neutrophil extracellular traps kill bacteria. Science 303, 1532–1535 (2004).

9. Wang, H. et al. Neutrophil extracellular traps in homeostasis and disease. Sig Transduct Target Ther 9, 235 (2024).

10. Law, S. M. et al. Neutrophil extracellular traps are associated with airways inflammation and increased severity of lung disease in cystic fibrosis. ERJ Open Research 10, (2024).

11. Obermayer, A. et al. New Aspects on the Structure of Neutrophil Extracellular Traps from Chronic Obstructive Pulmonary Disease and In Vitro Generation. PLOS ONE 9, e97784 (2014).

12. Grabcanovic-Musija, F. et al. Neutrophil extracellular trap (NET) formation characterises stable and exacerbated COPD and correlates with airflow limitation. Respir Res 16, 59 (2015).

13. Dworski, R., Simon, H.-U., Hoskins, A. & Yousefi, S. Eosinophil and neutrophil extracellular DNA traps in human allergic asthmatic airways. J Allergy Clin Immunol 127, 1260–1266 (2011).

14. Radermecker, C. et al. Neutrophil extracellular traps infiltrate the lung airway, interstitial, and vascular compartments in severe COVID-19. J Exp Med 217, e20201012 (2020).

15. Narasaraju, T. et al. Excessive Neutrophils and Neutrophil Extracellular Traps Contribute to Acute Lung Injury of Influenza Pneumonitis. Am J Pathol 179, 199–210 (2011).

16. Pillai, P. S. et al. Mx1 reveals innate pathways to antiviral resistance and lethal influenza disease. Science 352, 463–466 (2016).

17. Funchal, G. A. et al. Respiratory Syncytial Virus Fusion Protein Promotes TLR-4–Dependent Neutrophil Extracellular Trap Formation by Human Neutrophils. PLoS One 10, e0124082 (2015).

18. Cortjens, B. et al. Neutrophil extracellular traps cause airway obstruction during respiratory syncytial virus disease. J Pathol 238, 401–411 (2016).

19. Boboltz, A., Yang, S. & Duncan, G. A. Engineering *in vitro* models of cystic fibrosis lung disease using neutrophil extracellular trap inspired biomaterials. J. Mater. Chem. B 11, 9419–9430 (2023).

20. Linssen, R. S. et al. Neutrophil Extracellular Traps Increase Airway Mucus Viscoelasticity and Slow Mucus Particle Transit. Am J Respir Cell Mol Biol 64, 69–78 (2021).

21. Duncan, G. A. et al. Microstructural alterations of sputum in cystic fibrosis lung disease. JCI Insight 1, (2016).

22. Bornhöfft, K. F. et al. Sialylated Cervical Mucins Inhibit the Activation of Neutrophils to Form Neutrophil Extracellular Traps in Bovine in vitro Model. Front Immunol 10, 2478 (2019).

23. Mohanty, T. et al. A novel mechanism for NETosis provides antimicrobial defense at the oral mucosa. Blood 126, 2128–2137 (2015).

24. Val, S. et al. MUC5B induces in vitro neutrophil extracellular trap formation: Implication in otitis media. Laryngoscope Investig Otolaryngol 5, 536–545 (2020).

25. Gipson, I. K. et al. The Amount of MUC5B mucin in cervical mucus peaks at midcycle. J Clin Endocrinol Metab 86, 594–600 (2001).

26. Walters, M. S. et al. Generation of a human airway epithelium derived basal cell line with multipotent differentiation capacity. Respir Res 14, 135 (2013).

27. Birnie, G. D. The HL60 cell line: a model system for studying human myeloid cell differentiation. Br J Cancer Suppl 9, 41–45 (1988).

28. Delaveris, C. S. et al. Synthetic Siglec-9 Agonists Inhibit Neutrophil Activation Associated with COVID-19. ACS Cent. Sci. 7, 650–657 (2021).

29. Dömer, D., Walther, T., Möller, S., Behnen, M. & Laskay, T. Neutrophil Extracellular Traps Activate Proinflammatory Functions of Human Neutrophils. Front. Immunol. 12, (2021).

30. Brannon, E. R. et al. Polymerized Salicylic Acid Microparticles Reduce the Progression and Formation of Human Neutrophil Extracellular Traps (NET)s. Advanced Healthcare Materials 14, 2400443 (2025).

31. Jiang, D., Saffarzadeh, M. & Scharffetter-Kochanek, K. <em>In vitro</em> Demonstration and Quantification of Neutrophil Extracellular Trap Formation. Bio-protocol 7, (2017).

32. Tatsiy, O. & McDonald, P. P. Physiological Stimuli Induce PAD4-Dependent, ROS-Independent NETosis, With Early and Late Events Controlled by Discrete Signaling Pathways. Front Immunol 9, 2036 (2018).

33. Raskovalova, T. et al. Flow cytometric analysis of neutrophil myeloperoxidase expression in peripheral blood for ruling out myelodysplastic syndromes: a diagnostic accuracy study. Haematologica 104, 2382–2390 (2019).

34. McKenna, E. et al. Neutrophils: Need for Standardized Nomenclature. Front. Immunol. 12, (2021).

35. Doster, R. S., Rogers, L. M., Gaddy, J. A. & Aronoff, D. M. Macrophage Extracellular Traps: A Scoping Review. J Innate Immun 10, 3–13 (2018).

36. Song, D. et al. MUC5B mobilizes and MUC5AC spatially aligns mucociliary transport on human airway epithelium. Science Advances 8, eabq5049 (2022).

37. Harris, E. S. et al. Reduced sialylation of airway mucin impairs mucus transport by altering the biophysical properties of mucin. Sci Rep 14, 16568 (2024).

38. Kaler, L. et al. Mucus Physically Restricts Influenza A Viral Particle Access to the Epithelium. Advanced Biology 9, 2400329 (2025).

39. Lai, S. K., Wang, Y.-Y., Wirtz, D. & Hanes, J. Micro- and macrorheology of mucus. Adv Drug Deliv Rev 61, 86–100 (2009).

40. Abaricia, J. O., Shah, A. H. & Olivares-Navarrete, R. Substrate Stiffness Induces Neutrophil Extracellular Traps (NETs) Formation through Focal Adhesion Kinase Activation. Biomaterials 271, 120715 (2021).

41. Erpenbeck, L. et al. Effect of Adhesion and Substrate Elasticity on Neutrophil Extracellular Trap Formation. Front. Immunol. 10, (2019).

42. Joyner, K., Song, D., Hawkins, R. F., Silcott, R. D. & Duncan, G. A. A rational approach to form disulfide linked mucin hydrogels. Soft Matter 15, 9632–9639 (2019).

43. Suk, J. S. et al. Rapid transport of muco-inert nanoparticles in cystic fibrosis sputum treated with N-acetyl cysteine. Nanomedicine (Lond) 6, 365–375 (2011).

44. Du, W. et al. Mechano-induced patterned domain formation by monocytes. Nat. Mater. 25, 523–536 (2026).

45. Morawiec, M.-L. et al. Neutrophil extracellular traps in diseases of the female reproductive organs. Front Immunol 16, 1589329 (2025).

46. Crocker, P. R., Paulson, J. C. & Varki, A. Siglecs and their roles in the immune system. Nat Rev Immunol 7, 255–266 (2007).

47. Khatua, B., Bhattacharya, K. & Mandal, C. Sialoglycoproteins adsorbed by Pseudomonas aeruginosa facilitate their survival by impeding neutrophil extracellular trap through siglec-9. J Leukoc Biol. 91, 641–655 (2012).

48. Yu, H. et al. Siglec-8 and Siglec-9 binding specificities and endogenous airway ligand distributions and properties. Glycobiology 27, 657–668 (2017).

49. Ge, L., Wang, N., Chen, Z., Xu, S. & Zhou, L. Expression of Siglec-9 in peripheral blood neutrophils was increased and associated with disease severity in patients with AECOPD. Cytokine 177, 156558 (2024).

50. Welsh, K. G. et al. MUC5AC and a Glycosylated Variant of MUC5B Alter Mucin Composition in Children With Acute Asthma. Chest 152, 771–779 (2017).

51. Kirkham, S. et al. MUC5B Is the Major Mucin in the Gel Phase of Sputum in Chronic Obstructive Pulmonary Disease. Am J Respir Crit Care Med 178, 1033–1039 (2008).

52. Jeffries, J. L. et al. Pseudomonas aeruginosa pyocyanin modulates mucin glycosylation with sialyl-Lewisx to increase binding to airway epithelial cells. Mucosal Immunol 9, 1039–1050 (2016).

53. Liegeois, M. A. et al. Cellular and molecular features of asthma mucus plugs provide clues about their formation and persistence. J Clin Invest 135, e186889.

54. Newburger, P., Chovaniec, M., Greenberger, J. & Cohen, H. Functional changes in human leukemic cell line HL-60. A model for myeloid differentiation. J Cell Biol 82, 315–322 (1979).

55. Crowther, R. S. & Wetmore, R. F. Fluorometric assay of O-linked glycoproteins by reaction with 2-cyanoacetamide. Analytical Biochemistry 163, 170–174 (1987).

56. Müller, H. E. Occurrence of Neuraminidase and Acylneuraminate Pyruvate Lyase in a Strain of Vibrio Falling into Heiberg’s Group II Isolated from a Patient with Diarrhea. Infect Immun 8, 430–433 (1973).

57. Kaler, L. et al. Influenza A virus diffusion through mucus gel networks. Commun Biol 5, 249 (2022).

58. Schuster, B. S., Ensign, L. M., Allan, D. B., Suk, J. S. & Hanes, J. Particle tracking in drug and gene delivery research: State-of-the-art applications and methods. Advanced Drug Delivery Reviews 91, 70–91 (2015).

59. Schuster, B. S. et al. Overcoming the cystic fibrosis sputum barrier to leading adenoassociated virus gene therapy vectors. Molecular therapy : the journal of the American Society of Gene Therapy 22, 1484–93 (2014).

