## Supplemental Information for "Airway mucins function as endogenous inhibitors of neutrophil extracellular traps"

**Figure S1.** Histograms generated from flow cytometry data showing the fluorescence of a primary CD-14 antibody conjugated to PE-Cy7 in the isolated PBMC fraction versus PMN fraction.

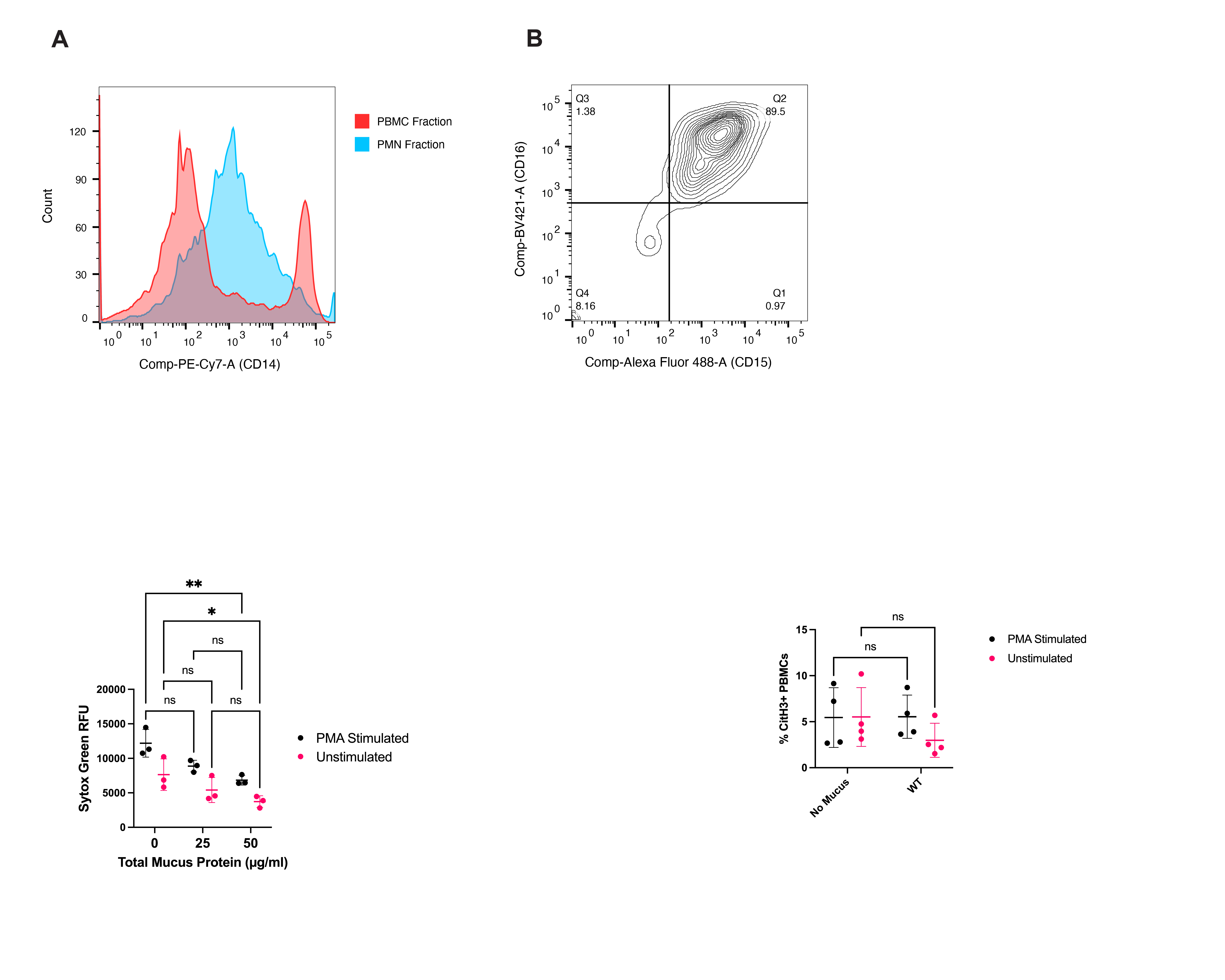

**Figure S2.** The gating strategy and representative flow cytometry plots for PMA stimulated PMNs with no mucus treatment compared to PMNs treated with WT HAE mucus at a final concentration of 50µg/ml total mucus protein.

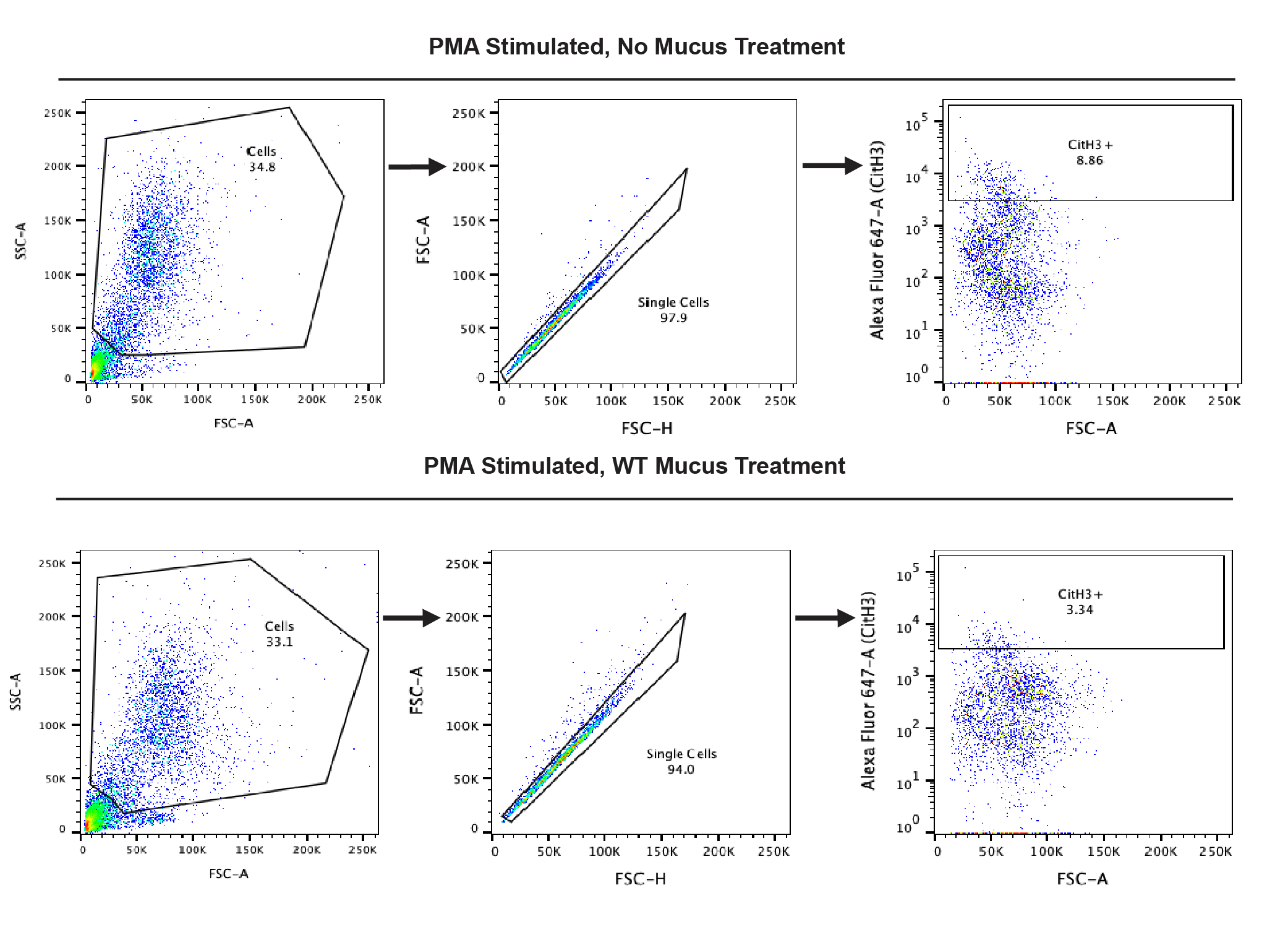

**Figure S3.** The isolated PBMC fraction was treated with WT mucus (50 µg/ml total mucus protein) or no mucus and either stimulated with PMA or left unstimulated (vehicle treatment). The cells were stained for CitH3 expression and the percentage of CitH3^+^ events was quantified using flow cytometry. Ns = not significant. Statistical test performed is a two way ANOVA with Šidák’s multiple comparisons test.

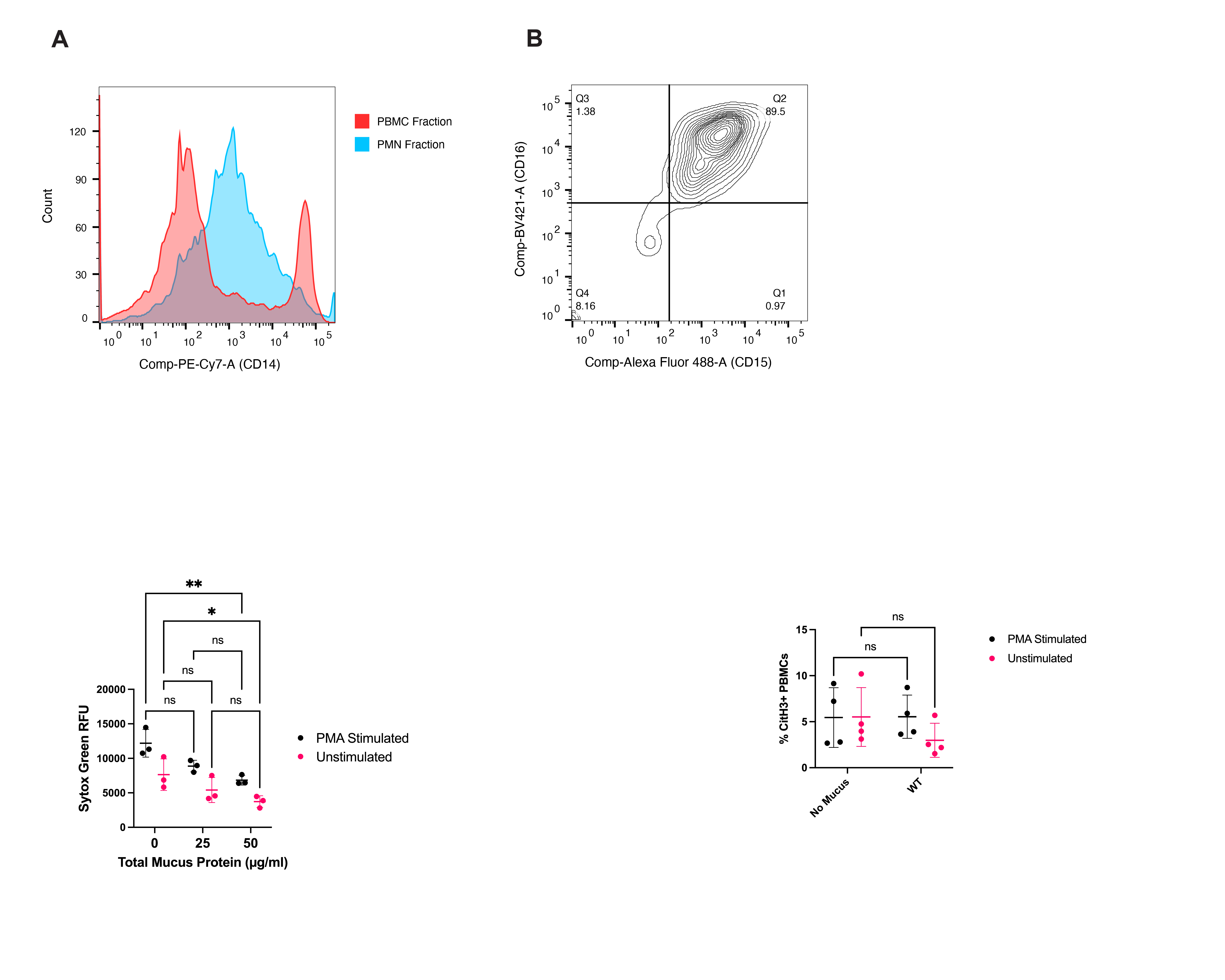

**Figure S4.** DHL-60 cells were treated with mucus (25 µg/ml total mucus protein) isolated from primary HAE cultures (primary) from healthy human donors or BCI-NS1.1 HAE cells (BCI) or received no mucus treatment. The dHL-60 cells were stimulated to undergo NETosis using PMA or left unstimulated and the Sytox green fluorescence was quantified. * p < 0.05, ** p < 0.01, ns = not significant. Statistical test performed is a two way ANOVA with Šidák’s multiple comparisons test.

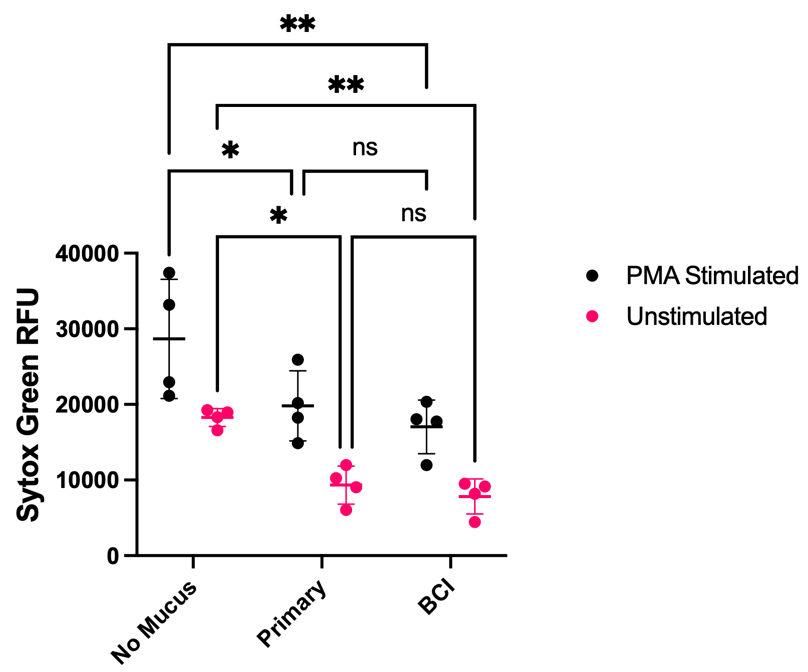

**Table S1.** Comparison of the fold changes in the NETosis markers of extracellular DNA (detected with Sytox green fluorescence) and CitH3. The fold change in each marker was calculated for each mucus treatment group in relation to the no mucus controls.

| Mucus Type | Fold Change in Sytox Fluorescence (Compared to No Mucus Control) | Fold Change in % CitH3^+^ (Compared to No Mucus Control) |
| --- | --- | --- |
| WT | 0.375 | 0.479 |
| AC KO | 0.767 | 0.857 |
| B KO | 0.739 | 0.681 |

**Figure S5.** Images of dHL-60 cells stimulated with PMA for 4 hours and stained for CitH3 expression. White arrows point to some of the dHL-60 cells undergoing NETosis. Scale bars, 20 µm.

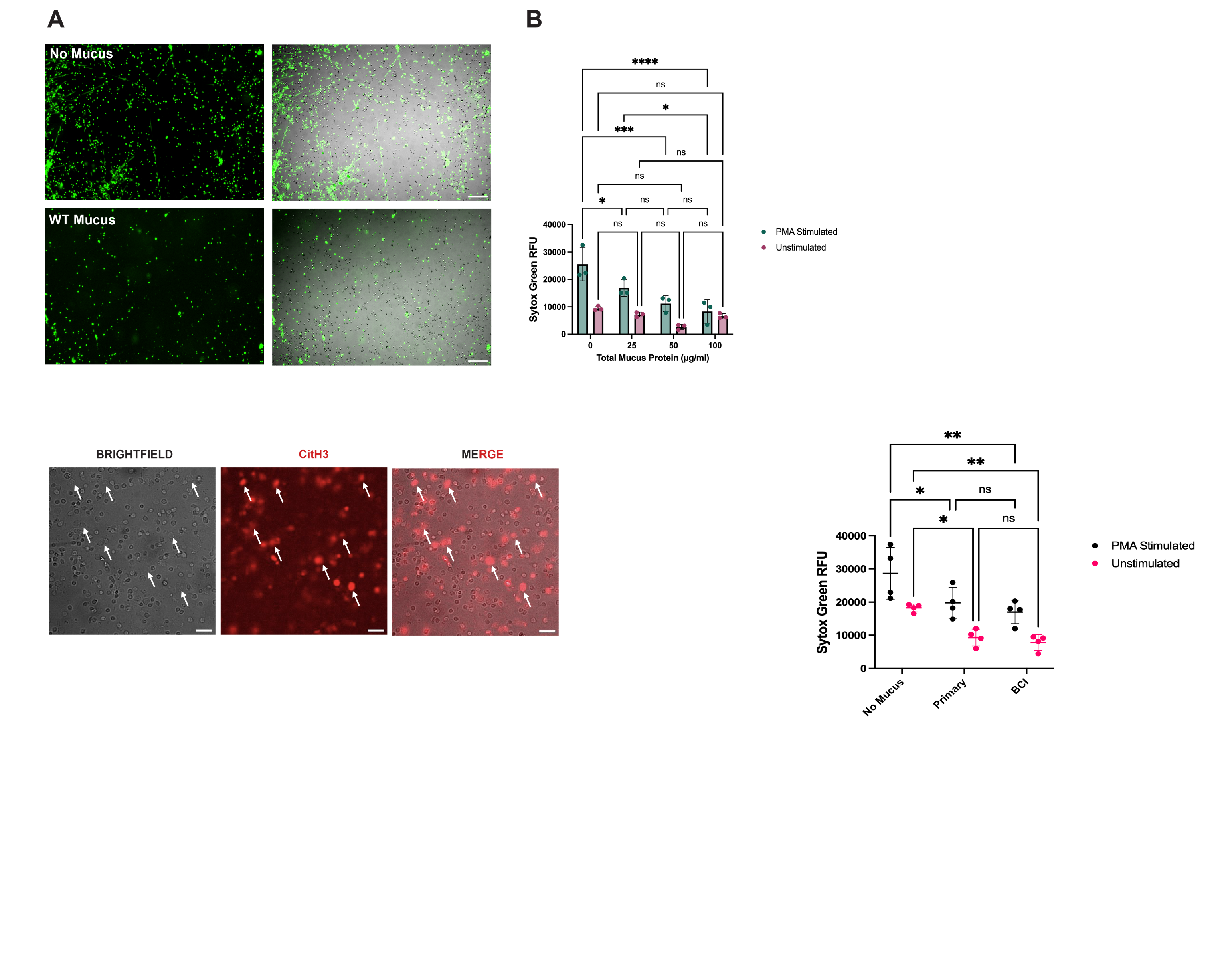

**Figure S6.** WT mucus was treated with a sialidase enzyme or a vehicle (mock) and A) the concentration of sialic acid was quantified in each sample. All samples were diluted to 100 µg/ml total mucus protein prior to measuring the sialic acid concentration. B) Sytox green fluorescence after treatment of dHL-60 cells with the mock or sialidase treated WT mucus (25 µg/ml total mucus protein final concentration) compared to no mucus treatment. C) WT mucus samples were subject to mock or sialidase treatment, but the cleaved glycans were retained in the sample and not removed using a de-salting column. Sytox green fluorescence of dHL-60 cells exposed to either mucus sample (final concentration of 25 µg/ml total mucus protein) compared to cells not treated with mucus. Cells were stimulated with PMA to undergo NETosis or left unstimulated. * p < 0.05, ** p < 0.01, *** p < 0.001, **** p < 0.0001, ns = not significant. Statistcal test performed in A) is an unpaired T-test and in B) and C) are two way ANOVAs with Šidák’s multiple comparisons test.

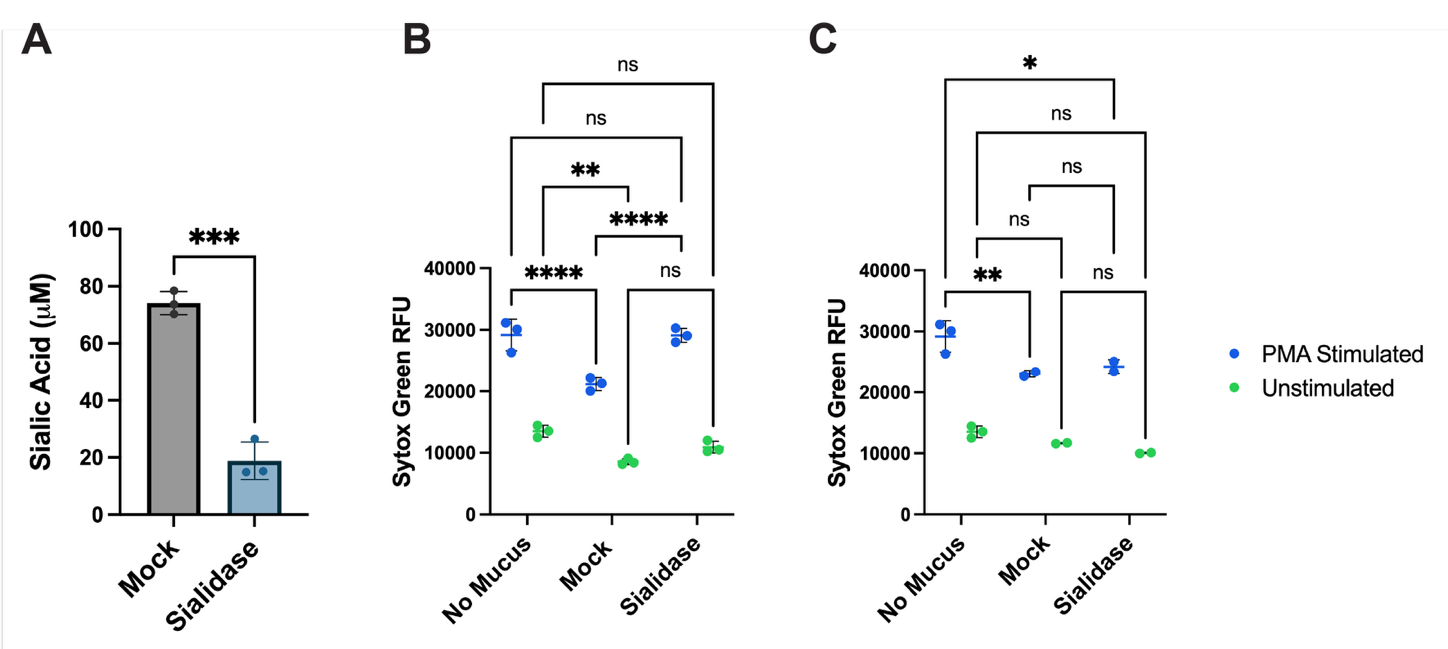

**Figure S7.** Sytox green fluorescence of dHL-60 cells treated with either porcine gastric mucin (PGM) or bovine submaxillary gland mucin (BSM) or no mucus. Each mucin type was dissolved in PBS at a concentration of 1 mg/ml and sterile filtered using a 0.45 µm pore size filter. PGM or BSM was added at a final concentration of 0.5 mg/ml to the media of the dHL-60 cells, the same concentration as what was incorporated into the hydrogel formulations. ** p < 0.01, *** p < 0.001, ns = not significant. Statistical test performed is a two way ANOVA with Šidák’s multiple comparisons test.

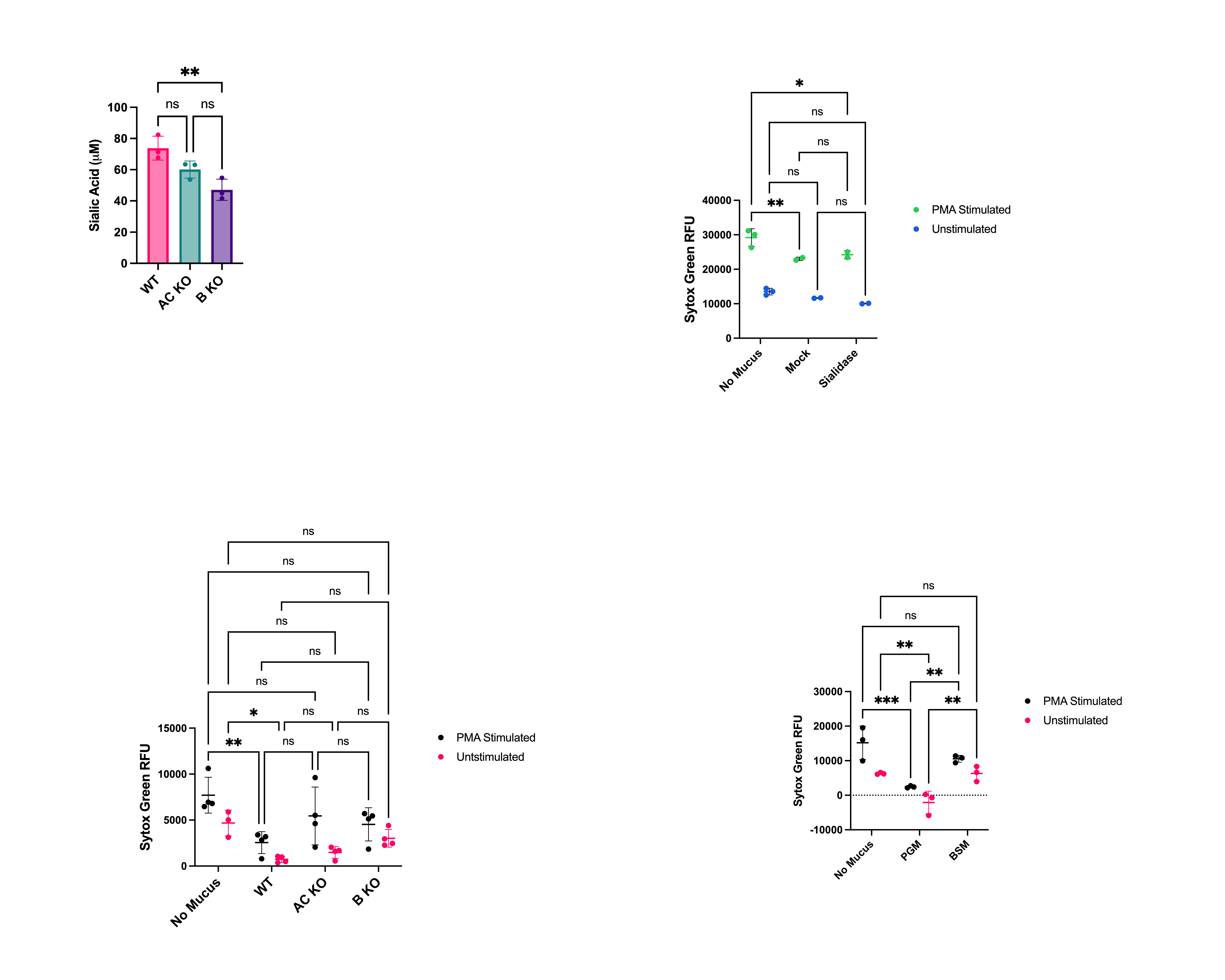
